# Disruption of the coordination between host circadian rhythms and malaria parasite development alters the duration of the intraerythrocytic cycle

**DOI:** 10.1101/791046

**Authors:** Amit K. Subudhi, Aidan J. O’Donnell, Abhinay Ramaprasad, Hussein M. Abkallo, Abhinav Kaushik, Hifzur R. Ansari, Alyaa M Abdel-Haleem, Fathia Ben Rached, Osamu Kaneko, Richard Culleton, Sarah E. Reece, Arnab Pain

## Abstract

Malaria parasites complete their intra-erythrocytic developmental cycle (IDC) in multiples of 24 hours (depending on the species), suggesting a circadian basis to the asexual cell cycle, but the mechanism controlling this periodicity is unknown. Combining *in vivo* and *in vitro* approaches using rodent and human malaria parasites, we reveal that: (i) 57% of *Plasmodium chabaudi* genes exhibit 24 h “circadian” periodicity in transcription; (ii) 58% of these genes lose transcriptional rhythmicity when the IDC is out-of-synchrony with host rhythms; (iii) 9% of *Plasmodium falciparum* genes show circadian transcription under free-running conditions; (iv) Serpentine receptor 10 (SR10) has a circadian transcription profile and disrupting it in rodent malaria parasites shortens the IDC by 2-3 hours; (v) Multiple processes including DNA replication and the ubiquitin and proteasome pathways are affected by loss of coordination with host rhythms and by disruption of SR10. Our results show that malaria parasites are at least partly responsible for scheduling their IDCs explaining the fitness benefits of coordination with host rhythms.

---

Daily fluctuations in environmental parameters such as light and temperature are assumed to select for the evolution of circadian clocks. In many organisms, circadian clocks schedule rhythms in behaviour, physiology and metabolism in coordination with periodic oscillations in the external environment^1^. The components of circadian clocks are diverse^2–4^. Many clocks function following the principles of transcription-translation feedback loops (TTFLs); a ‘clock’ protein inhibits the transcription of the gene that encodes it^5^ and clocks that operate via non-transcriptional oscillators^6^ and post-transcriptional control have also been identified^7^.

Many parasite species exhibit daily rhythms in behaviour and/or development that are scheduled to optimally exploit periodicities in transmission opportunities and/or resource availability^8, 9^. The parasitic protozoan *Trypanosoma brucei,* for example, possesses an intrinsic circadian clock that drives metabolic rhythms^10^. Malaria parasites complete their intra-erythrocytic developmental cycle (IDC) in 24h (or multiples of 24 h depending on the parasite species), suggesting a circadian basis to the asexual cell cycle^11^.

Rhythms in host feeding and innate immune responses influence the timing of rhythms in the IDC of rodent malaria parasites^12, 13^. Specifically, completion of the IDC, a glucose-demanding process, coincides with host food intake, and quiescence during the early phase of the IDC coincides with the daily nadir in host blood glucose that is exacerbated by the energetic demands of immune responses^12^. However, the extent to which malaria parasites or their hosts are responsible for IDC scheduling is unclear^14^. Either parasites are able to respond to time-of-day cues provided by the host to organise when they transition between IDC stages and complete schizogony, or parasites are intrinsically arrhythmic and allow the host to impose rhythms on the IDC (for example, restricting access to a nutrient that is essential to a particular IDC stage to a certain period each day, starves or kills parasites following the wrong IDC schedule).

Establishing how the timing and synchronicity of the IDC is established is important as temporal coordination with host rhythms is beneficial for parasite fitness^15, 16^, and because tolerance to antimalarial drugs is conferred to parasites that pause their IDCs^17–19^. Here, we use a combination of rodent malaria parasites *in vivo* and human malaria parasites *in vitro* to investigate the relationship between the IDC and host circadian rhythms.

Firstly, we identify components of the *P. chabaudi* transcriptome with 24 h periodicities and determine what happens to them, including the downstream biological processes, when coordination with host rhythms is disrupted (i.e. when the parasites’ IDC is “out of phase” with the host). Secondly, we show that *P. falciparum* also has a transcriptome with 24 h periodicity, even in the absence of host rhythms. Thirdly, we identify a transmembrane serpentine receptor with circadian expression in both species and demonstrate it plays a role in the duration of the IDC. Loss of this serpentine receptor disrupts many of the same processes affected when coordination to host rhythms is perturbed. Taken together, our results imply that malaria parasites are, at least in part, able to control the schedule of their IDCs. Coordinating the IDC with host rhythms could allow malaria parasites to enhance their fitness by facilitating temporal compartmentalisation of the IDC processes with respect to each other, by maximizing exploitation of resources provided by the host in a rhythmic manner, and by protecting vulnerable IDC stages from encountering peaks in the rhythms of host immune defenses.

## RESULTS

### The transcriptome of *Plasmodium chabaudi* responds to host circadian rhythms

Transcriptome analyses of time series RNA sequencing datasets were performed with *P.chabaudi* parasites from infections that were in synchrony (phase aligned; “host rhythm matched”) and out of synchrony (out of phase; “host rhythm mismatched”) with host circadian rhythms (for details see methods section). Briefly, a controlled number of infected red blood cells (RBC) were sub-inoculated from donor mice into two groups of recipient mice (n = 43 per group) each housed under different light regimes (12 hours (h) difference in lights on/off). In one group, the light regime was maintained as for the donor mouse (i.e. host rhythm matched, lights on: 7.30 (ZT0/24 (Zeitgeber Time: hours after lights on)) and lights off: 19.30(ZT12)). In the other group, the light-dark cycle was reversed (host rhythm mismatched, lights on: 19.30 (ZT0/24), lights off: 7.30 (ZT12)) (**Fig. 1a**). The effect of mismatch to host rhythms on the parasite was assessed by analyzing the parasite transcriptome every 3 h for 30 h (n=4 mice/group/time point).

**Fig. 1.**
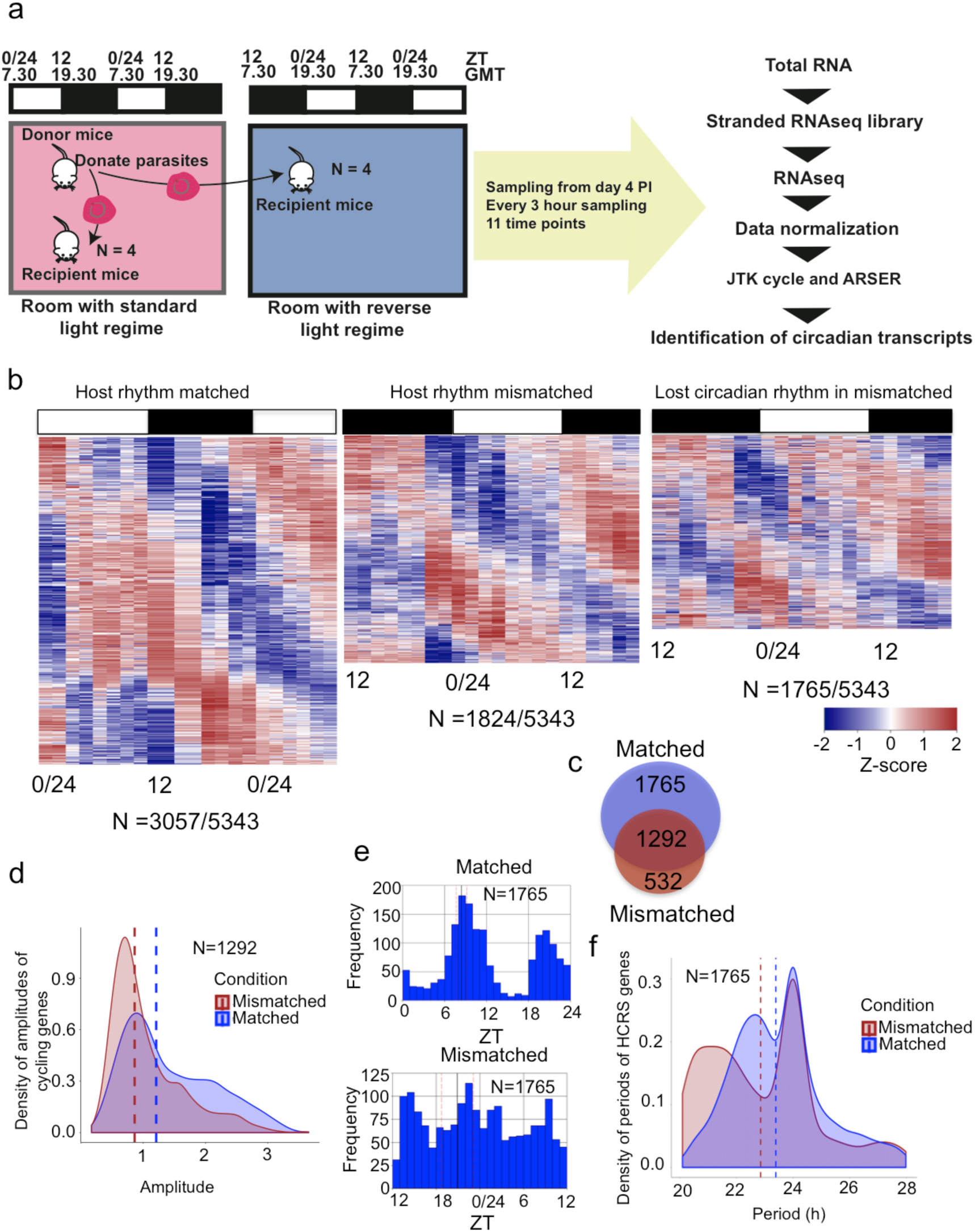
The transcriptome of *Plasmodium chabaudi* is sensitive to the phase of host circadian rhythms. **a)** Ring stage parasites from donor mice housed at a standard light regime were used to infect recipient mice housed in two rooms that differed by 12 hours in their light:dark cycle. Blood samples were collected for RNAseq analysis from day 4 post-infection every 3 hours for 11 time points (N = 4 mice per group per time point). ZT is Zeitgeber Time: hours after lights on. **b)** Time series gene expression heatmap views of transcripts with 24 hour (putative “circadian”) rhythmicity in matched and mismatched parasites. Right most side heatmap shows the expression pattern of 24 h transcripts in mismatched parasites that lost rhythmicity. Each row in each heatmap represents a single gene, sorted according to the phase of maximum expression starting from first sample time point. The phase of expression of each gene was obtained from ARSER output and N represents number of genes identified by both JTK^74^ and ARSER^75^ as fluctuating in expression in a 24 h manner. Each time point is represented by expression heatmap of two biological replicates. **c)** Venn diagram of number of 24 h (putative “circadian”) genes identified by both JTK and ARSER programs in matched (top) and mismatched (bottom) parasites. **d)** Transcripts with 24 h (putative “circadian”) rhythmicity in both matched and mismatched parasites had lower median amplitude (0.86, brown dashed line) in mismatched parasites compared to matched parasites (1.22, blue dashed line). **e)** Phase distributions of genes that displayed 24 h (putative “circadian”) rhythmicity only in matched parasites in the form of a histogram. The mean circular phase is indicated by a solid black line. N represents the number of cycling transcripts. The pink line represents the standard deviation of the mean of phases. Whilst these genes are not identified as having 24 h rhythms in mismatched parasites, their distribution is shown for comparison. **f)** Transcripts that displayed 24 h (putative “circadian”) rhythmicity only in matched parasites have median period close to 24 h (blue dashed line) and (for comparison) 23 h in mismatched parasites (brown dotted line).

After mismatch to host circadian rhythms, the IDC of *P. chabaudi* becomes rescheduled within approximately seven cycles to match the host’s rhythms. By the time of sampling (days 4-5 post infection, PI), the parasites were six hours mismatched to the host’s rhythm (**Supplementary Fig. 1a**). Specifically, in mismatched parasites, merozoite egress from RBCs following schizogony peaked six hours after matched parasites (ZT 0/24 and ZT 18 respectively; inferred from ring stage rhythms). Parasites in both the matched and mismatched infections remained synchronous throughout the sampling period (**Supplementary Fig. 1a**).

After quantifying gene transcription at each time point through RNAseq analysis, we identified genes that followed ∼24 h rhythms in transcription according to two commonly used and independent algorithms (see Methods). Genes were considered to have putative circadian rhythms (hereafter referred to as “circadian”) if both algorithms independently detected a ∼24 h periodicity in their transcription with a threshold of *p* < 0.05. Of a total of 5,343 genes in *P. chabaudi* (5,158 detected and considered for analysis), 3,057 (58%) in matched parasites, and 1,824 (34%) in mismatched parasites, exhibited circadian rhythms in transcription (**Fig. 1b**, **Supplementary Data 1**). A permutation test was performed to empirically determine the false discovery rate (FDR) in detecting circadian transcripts. The original time points were shuffled randomly for 1,000 iterations and circadian transcripts were identified each time by both the algorithms. The permutation test identified that the number of circadian transcripts in both matched and mismatched parasites detected in the original sampling order by both the programs was significantly higher (FDR < 0.05, **Supplementary Fig. 1b**) than when the sampling order was randomly permuted indicating that the circadian genes identified were transcribed in a rhythmic manner above background noise.

### Changes in rhythmicity due to misalignment of parasite and host rhythms

Over 80% of the genes expressed during the IDC of malaria parasites undergo a tight temporal expression cascade associated with development into specific parasite stages^20, 21^. The periodicity of the transcription profile of these genes can be ∼ 24 h, 48 h or 72 h depending on the malaria parasite species: for *P. chabaudi,* the transcription profile of these genes have ∼ 24 h periodicity while for *P. falciparum* they have ∼48 h periodicity. This means that for *P. chabaudi,* genes that are transcribed in association with the IDC cascade and genes that encode proteins that could be involved in a circadian clock or its outputs all display 24 h periodicity in their transcription profiles. However, IDC genes should follow transcription rhythms that peak six hours later in mismatched compared to matched parasites, whereas gene transcripts that follow the phased rhythm in both mismatched and matched parasites with respect to host ZT may be involved in a circadian clock or its outputs. Genes that lose transcription rhythmicity in mismatched parasites may do so because the IDC and/or homeostasis are negatively affected by misalignment with host rhythm, as a consequence of re-aligning the IDC schedule with host circadian rhythms, or both.

Comparing rhythmic transcripts detected in both matched and mismatched parasites identified three sets of transcripts: 1) 1,765 genes (33% of the total genes) with 24 h rhythmic transcription (*p* < 0.05) in matched parasites that exhibit an arrhythmic transcription profile in mismatched parasites (*p* > 0.05); 2) 1,292 genes with transcription profiles with 24 h rhythm in both matched and mismatched parasites; and 3) 532 genes whose transcription profiles were arrhythmic in matched parasites and rhythmic in mismatched parasites (*p* < 0.05, Fig. 1b and 1c, **Supplementary Fig. 1c**). Hierarchical clustering analysis identified biological replicates to be tightly clustered (**Supplementary Fig. 1d**). Comparison of the 11 time points using principal component analysis identified the first component in both the conditions with a cyclic pattern that accounted for > 85 % of total variance (**Supplementary Fig. 1e**), which supports the hypothesis that the large number of transcripts detected as circadian are truly transcribed at a 24 h periodicity.

Out of 1,292 genes that were rhythmically transcribed in both matched and mismatched parasites, we found 685 genes (53%) that had a delayed phase of transcription with a delay of about 6 h (± 1.5 h) in mismatched compared to matched parasites (**Supplementary Fig. 1f**) which complements our phenotypic observation of the phase difference between mismatched and matched parasite IDCs (**Supplementary Fig. 1a**). Gene ontology enrichment analysis revealed that biological processes associated with these transcripts include DNA metabolic processes and cellular responses to stress (FDR < 0.05). The other 607 genes display broad differences in their phase of transcription between matched and mismatched parasites, and so might also be IDC associated genes affected by a loss of scheduling forces and / or genes undergoing readjustment of their phase of transcription to realign the IDC with the host rhythm. The amplitude of rhythmically transcribed genes is significantly higher (*p* < 0.0001, unpaired student *t* test) in matched compared to mismatched parasites (**Fig. 1d**), suggesting a loss of synchronicity in the transcription of mismatched parasites that is not severe enough to impact on IDC synchronicity as measured by stage proportions (**Supplementary Fig. 1a**). Such a dampening of rhythms is a typical consequence of misalignment of a circadian clock with its time-of-day cue^22^.

Arrhythmically transcribed genes (N=532) in matched parasites that exhibit rhythmic transcription in mismatched parasites enriched to a single gene ontology biological process term; RNA processing/splicing. Why genes should gain rhythmicity of transcription in mismatched parasites is unclear; it is possible that they compensate for stresses imposed by being misaligned to host rhythms or they may represent an alternative set of IDC genes that the parasite expresses during its rescheduling period. Such a phenomenon is proposed to operate in humans, by which a set of transcripts gains rhythmicity in older individuals coinciding with the loss of canonical clock function^23^.

The most striking observation is that 1,765 genes (33%) lose rhythmicity of transcription in mismatched parasites. In matched parasites, a bimodal distribution of transcription for these genes was observed with peaks at two different times of the day (ZT 8 and ZT 20, **Fig. 1e**) corresponding to the late trophozoite and ring stages of the IDC, respectively. Whilst these genes have a periodicity of transcription extremely close to 24 h in matched parasites (median periodicity =23.89h), 55% of these transcripts had shorter periodicities in mismatched parasites (between 20h to 24h), which in turn reduced the overall periodicity by ∼ 1 hour (median periodicity 22.85h) (**Fig. 1f**). Period estimates from genes that lost rhythms in transcription profiles are used here to illustrate an overall trend, rather than provide information on individual genes. It is possible that the shorter periods for mismatched parasites correlates with a shorter IDC, thus explaining how mismatched parasites become rescheduled by approx. six hours within four cycles of replication (an average of 1.5 h per cycle).

We undertook further analysis of the 1,765 genes that lose rhythmicity in their transcription profile in mismatched parasites to examine which biological processes are affected. We divided genes into 12 groups based on the time of day (“phase”) of their maximal transcription and performed gene ontology (GO) based enrichment analysis within each group every two hours (using the fitted model output from ARSER). The point at which there were the highest relative copy numbers of mRNA from these genes present in samples was taken to be indicative of an overall trend. A wide range of biological processes including carbohydrate metabolism, nucleotide and amino acid metabolism, DNA replication, oxidation-reduction processes, translation, RNA transport, aminoacyl-tRNA biosynthesis and ubiquitin mediated proteolysis and proteasome pathways were enriched in different phase clusters and so appear to be under the influence of host rhythms (**Fig. 2a** **and Supplementary Data 2**). Many of these biological processes are under circadian clock control in other organisms^24, 25^. Disruption to any (or all) of these processes could explain the 50% reduction in parasite densities observed for mismatched parasites by O’Donnell et. al.^15, 16^. To gain insight into these perturbed processes, we assessed these findings on a gene-by-gene basis in the context of the IDC and how it might be scheduled.

**Fig. 2.**
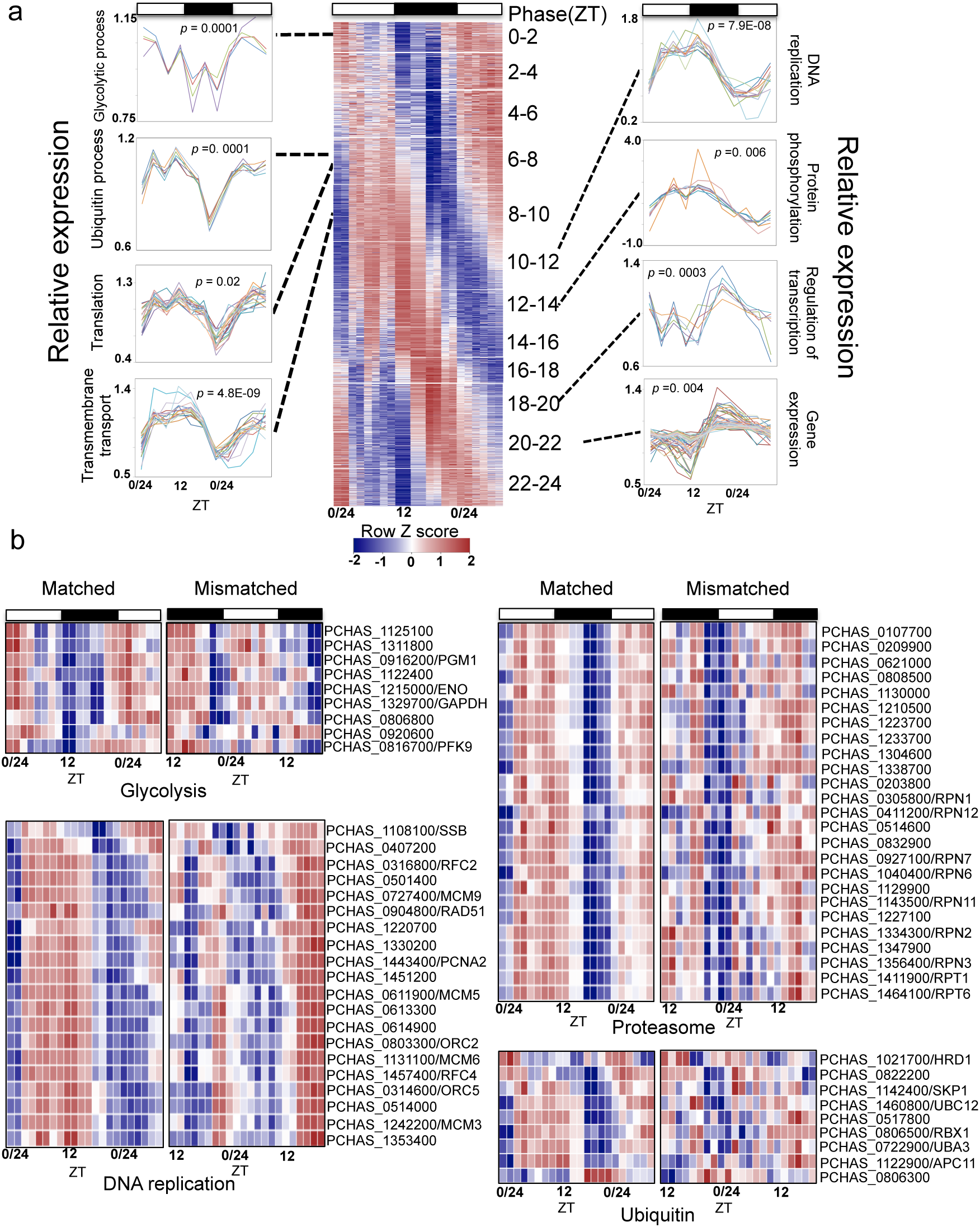
Multiple key biological pathways are affected by mismatch to the phase of the host’s circadian rhythm in the rodent malaria parasite *Plasmodium chabaudi.* **a)** Time series gene expression view of genes that displayed with 24 h (putative “circadian”) rhythmicity in matched parasites but lost rhythmicity in mismatched parasites. Genes were sorted based on phase of maximum expression and segregated into 12 groups with each group representing 2 hours (h) phase clusters. Line plots along the sides of the heat map represent expression profiles of individual genes significantly enriched to gene ontology terms (*p*<0.05, hypergeometric test) representing few crucial biological processes. Each plot has information about the false discovery rate corrected *p* value of representing GO term. The Y axis represents relative expression of genes at each time point determined by count level expression of each gene normalized by its mean across 11 time points. **b)** Heatmaps illustrating the expression patterns of circadian genes for matched and mismatched parasites that are involved in the ubiquitin and proteasome systems, and the DNA replication and glycolysis pathways. These genes lost circadian rhythmicity in mismatched parasites. Genes have been sorted based on the phase of maximum expression. The colour scheme represents the row Z score. Each time point is represented by the expression heatmap of two biological replicates.

### Energy metabolism

Blood glucose levels are rhythmic during the circadian cycle in mice^26^. This rhythm is exacerbated in malaria-infected mice due to the alterations in energy metabolism experienced by inflammatory leukocytes^12^. Given that completing the IDC is glucose demanding, malaria parasites may be expected to express genes rhythmically to utilize the energy source efficiently. This could be achieved via the recently discovered nutrient sensing mechanism and that allows parasites to respond to alterations in glucose availability through transcriptional rearrangement^27^. In support of this, genes involved in energy metabolism pathways (glycolysis, fructose and mannose metabolism) showed circadian transcription patterns in the matched parasites while this rhythmicity was lost in mismatched parasites (**Figs. 2a and 2b**, **Supplementary Data 2**). The maximum relative number of transcripts of glycolysis-associated genes in matched *P. chabaudi* was observed between ZT 22-ZT 2, which corresponds to the ring and early trophozoite stages of the parasite and are also when genes associated with glycolysis are maximally transcribed in *P. falciparum in vitro*^20^. Genes associated with glycolysis pathways in *Trypanosoma brucei* are under the control of an endogenous circadian clo^10^. If malaria parasites also possess such an oscillator, or another controller of the IDC, misalignment to the host rhythm should disrupt glucose metabolism.

### The ubiquitin-proteasome system

In chronobiology, the ubiquitin-proteasome system (UPS) plays a direct role in determining the half-life of core clock components and other clock controlled protein functions^28^. The conserved UPS system works by tagging ubiquitin (Ub) to a protein destined for enzymatic degradation and the tagged proteins are then recognized by a multi-catalytic protease complex called the 26S proteasome system that eventually degrades the tagged proteins. A total of nine out of 25 (36%) genes associated with the ubiquitin mediated proteolysis pathway and 25 out of 32 (78%) genes encoding core and regulatory components of the proteasome system lost transcriptional rhythmicity in mismatched parasites (**Fig. 2b**, **Supplementary Data 2**). Notably, genes that lost transcriptional rhythmicity include those associated with one E1 Ub-activating enzyme, four E2 Ub-conjugating enzymes, and three ring finger type E3 Ub-ligases (RBX1, SYVN and Apc11) from the ubiquitin mediated proteolysis system. Furthermore, rhythmicity was also lost in RBX1, Apc11 and the adaptor protein SKP1 from the proteasome pathway. These genes are part of the anaphase promoting complex (APC) which is a cell cycle regulated ubiquitin protein ligase. Mutation studies in budding yeast show that *rbx1*and *apc11* are essential for APC activity^29–32^.

The majority of ubiquitin-proteasome associated genes have a peak of transcription between ZT 7.5-10 in matched parasites, which corresponds to the late trophozoite/ schizont stage of the IDC (**Fig. 2b**). The UPS plays a crucial role for the liver, blood and sexual stages of the parasite. In the blood stage, particularly during the late IDC stage, the UPS helps parasite to shift from its involvement in generic metabolic and cellular machinery to specialized parasite function. Loss of rhythms of UPS associated genes suggests that part of the UPS system that plays a role in regulating important events in cell division and in protein homeostasis is detrimentally affected by mismatch with the host rhythm.

### DNA replication associated genes

GO enrichment analysis revealed that 19 out of 43 (44%) genes associated with DNA replication lost transcriptional rhythmicity in mismatched parasites (**Figs. 2a and 2b and Supplementary Data 2**). These included genes encoding subunits of DNA polymerase, replication-licensing factors, DNA helicase and the DNA repair protein RAD51. Genes associated with DNA replication reached peak transcription between ZT8-12 in matched parasites, which corresponds to the transition from the late trophozoite to the schizont stage during the IDC, and is when DNA replication machinery components are transcribed^20^.

Other cell cycle associated genes encoding cdc2-related protein kinase 4 and 5, anaphase-promoting complex 4, cyclin 1, cullin like protein, regulator of initiation factor 2 and replication termination factor also lost transcriptional rhythmicity in mismatched parasites. Biological clocks control the timing of DNA replication in many organisms^33–37^. If the parasite uses a clock that follows host circadian rhythm to initiate its own DNA replication to remain in synch with host rhythms, then when the cue to begin DNA replication is received, mismatched parasites will be at the wrong IDC stage to achieve the task, resulting in disorganized DNA replication.

### The redox system

Organisms employ multiple antioxidant mechanisms such as the use of super oxide dismutase, peroxiredoxins, glutathione and thioredoxin systems to remove harmful reactive oxygen species radicals that include hydrogen peroxides and super oxide ions. Seven genes out of 31 associated with cell redox and glutathione metabolism lost transcriptional rhythmicity in mismatched parasites and are enriched for the term ‘cell redox homeostasis’ (*Padj* < 0.001; **Supplementary Data 2**). All seven genes displayed maximum transcription during ZT6-8 in matched parasites. Peroxiredoxin proteins show circadian rhythmicity in oxidation/reduction cycles and these are conserved across the tree of life^3^. Although genes that encode peroxiredoxins are not necessarily expressed in a circadian manner during circadian oxidation/reduction cycles^38^, two out of three genes encoding peroxiredoxin in *P. chabaudi* showed circadian transcription in both matched and mismatched parasites, while one gene (PCHAS_0511500) lost transcriptional rhythmicity in mismatched parasites. As for glycolysis, the expression of genes involved in redox metabolism is also driven by an endogenous clock in *T. brucei*^10^. It is possible that redox metabolism associated genes are also driven by a clock in malaria parasites and misalignment with host rhythms interferes with the parasites’ ability to prepare for and cope with redox challenges.

### 24-hour endogenous rhythms in the transcriptome of *Plasmodium falciparum*

We used *P. falciparum* to investigate whether malaria parasites schedule their IDC to coincide with host rhythms. Parasites may set the timing of IDC transitions using a circadian clock that is entrained by a host rhythm (i.e. a Zeitgeber). One of the criteria for demonstrating clock control of a rhythm is that the rhythm persists (“free-runs”) under constant conditions. *In vitro* culture can provide constant conditions and in contrast to *P. chabaudi, P. falciparum* has an IDC of approx. 48 h, allowing putative clock genes and their downstream interactors to be distinguished from IDC genes. Observing a rhythm of 24 h is consistent with the presence of a circadian clock, but other criteria such as temperature compensation and entrainment must be fulfilled to conclude the presence of a clock.

We analysed the published 48-hour time-series IDC microarray expression profile of *P. falciparum* captured at a 1 h time-scale resolution by^20^. Temperature is the Zeitgeber for trypanosome clocks^10^, but *P. falciparum* experienced a constant temperature in the study of Bozdech *et al.* (2003)^20^. Furthermore, Bozdech et al. (2003)^20^ cultured *P. falciparum* in constant darkness, though light:dark rhythms do not influence the schedule of the *P. chabaudi* IDC^13^. Expression data from two odd time-points are missing in this dataset so we considered only data from the even time-points, giving the dataset a 2 h resolution. We identified 494 transcripts (9% of the genes) with ∼24 h rhythmicity (*q* < 0.05) from the expression profiles of 4,818 genes (**Fig. 3a**, **Supplementary Data 3**). The median amplitude of oscillations of the 24 h free-running genes was 0.20, which is lower than the amplitude of the rhythmic genes detected in *P. chabaudi in vivo* (1.22). A lower amplitude of 24 h expression rhythms in *P. falciparum in vitro* is expected as it is devoid of exposure to host circadian rhythms, whereas a higher amplitude of 24 h expression rhythms in *P. chabaudi in vivo* may be maintained by exposure to host rhythms.

**Fig. 3.**
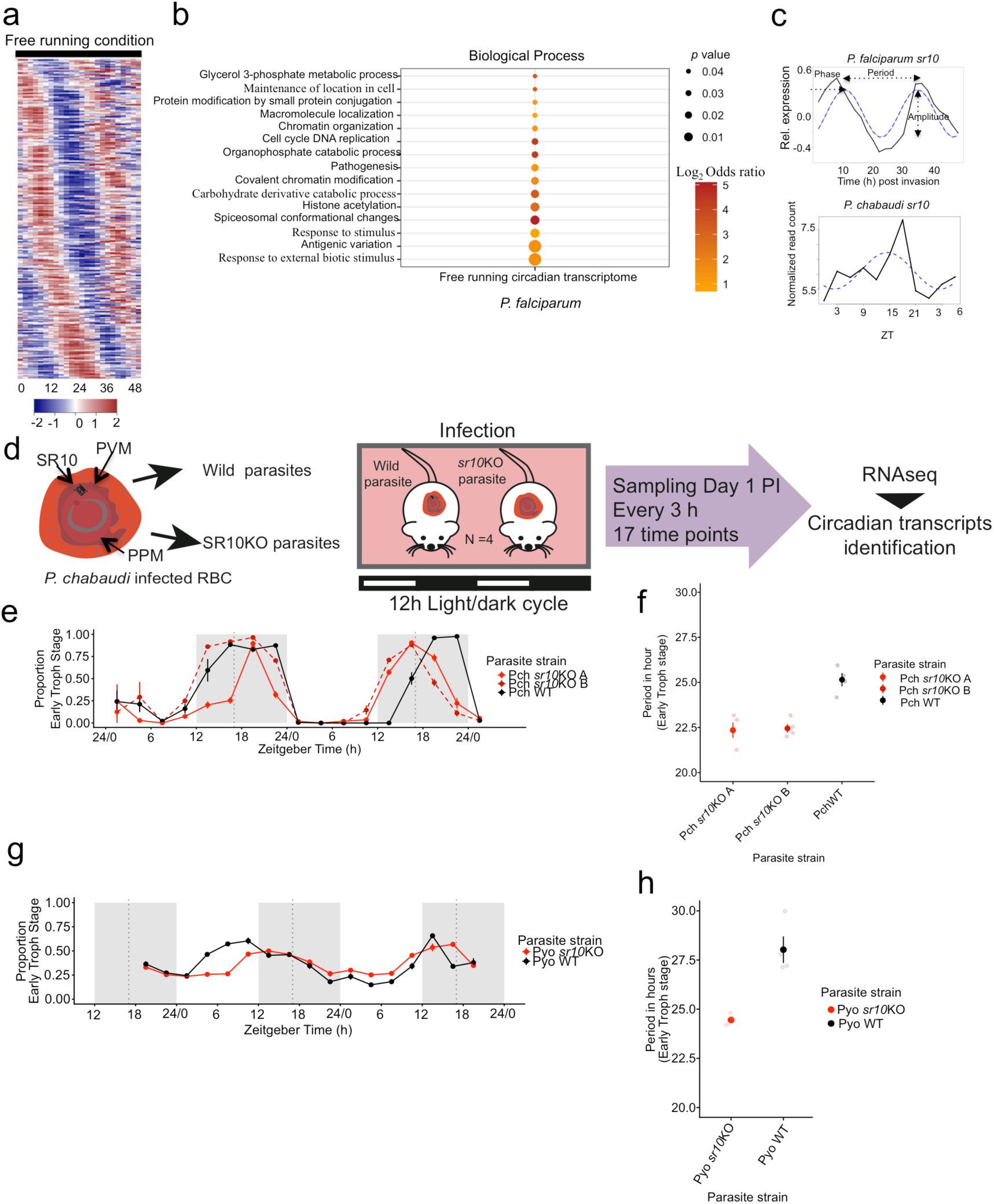
Expression of serpentine receptor 10 is “circadian” in *Plasmodium falciparum in vitro* and maintains the duration of the intra-erythrocytic developmental cycle of *P. chabaudi* and *P. yoelii in vivo*. **a)** Time series gene expression heat map view of circadian genes identified in *P. falciparum in vitro* in “free running” (constant) conditions. Genes sorted based on the phase of maximum expression starting from time T:0 post merozoite invasion. *P. falciparum* IDC expression data was obtained from Bozdech et al., (2003)^20^. **b)** Manually curated gene ontology terms enriched for *P. falciparum* genes with 24 h expression (*Padj* < 0.05). **c)** Line graphs represent the expression of serpentine receptor 10 in *P. falciparum* over its 48 h IDC (top plot) and in *P. chabaudi* during its 24 h IDC (bottom plot). Dotted lines show the best-fit sinusoidal curves. *P. falciparum* IDC expression data was obtained from^20^ and *P. chabaudi* expression data from this study. **d)** *P. chabaudi* wild-type and s*r10*KO parasites were used to initiate infections in CBS mice. Blood was collected from day 1 (ZT 13.5) every 3 h during the following 48 hrs. Expression data from two biological replicates over 14 time points (from day 2 PI, ZT22.5) were analyzed to identify “circadian” transcripts. Abbreviations: PVM, parasitophorous vacuole membrane; PPM, parasite plasma membrane; RBC, red blood cell. Proportion of parasites in the blood at early trophozoite stage in *P. chabaudi* wild type and *sr10*KO clones (Mean ± SEM, N=4/clone) **e)** IDC duration of *P. chabaudi* wild type and *sr10*KO clones (Mean ± SEM, N=4/clone). **f)** Proportion of parasites in the blood at early trophozoite stage in *P. yoelii* wild type and *sr10*KO clones (Mean ± SEM, N=4/clone). **g)** IDC duration of *P. yoelii* wild type and *sr10*KO clones (Mean ± SEM, N=4/clone).

Genes identified in *P. falciparum* with a circadian transcription profile are enriched for most of the processes that lost rhythmicity in mismatched *P. chabaudi* infections (carbohydrate metabolism, DNA replication and the ubiquitin proteasome system (**Fig. 3b**). In contrast to mismatched *P. chabaudi*, no gene transcripts related to redox processes were enriched. This suggests that if *P. falciparum* has a clock, it is not used to anticipate daily redox challenges and that instead redox genes are transcribed only in response to a change in redox levels inside the host (possibly as a consequence of host metabolism). Furthermore, this finding suggests that circadian rhythms in the redox state of RBCs^7^, which persist when RBCs are cultured in constant conditions, are either not important for malaria parasites or are dampened by the developing parasite.

We also identified individual members of different sub-telomeric multigene families that are transcribed in a circadian manner in both *P. falciparum* and *P. chabaudi* datasets (**Supplementary Data 3**). Multigene families whose members are present in the subtelomeric regions of the *P. falciparum* chromosomes^39^ and *P. chabaudi* chromosomes^40^, and have at least three paralogous members in the genome were considered for this analysis. These included members of major antigenically variant gene families such as *var*, *riffins* and other families such as (Maurer’s cleft two-transmembrane protein encoding genes and *Plasmodium* helical interspersed subtelomeric protein encoding genes). Further analysis of host circadian rhythm-matched and mismatched *P. chabaudi* transcriptome revealed circadian expression of a large subset of RMP (rodent malaria parasites) sub-telomeric multigene families such as *cir*, RMP-*fam-a* and *etramp* (early transcribed membrane protein), a large proportion of which lose their rhythmic expression pattern in mismatched parasites (**Supplementary Data 3**). The precise biological significance of the rhythmic expression of these genes needs detailed future experimentations.

Finally, comparison of the transcriptional profile of genes that lost rhythm in mismatched *P. chabaudi* (N =1,765) to orthologues of “free running” *P. falciparum* rhythmic genes (N = 404, one to one orthologues out of 495 genes) identified 110 common genes. The transcription of these genes is sensitive to the timing of host circadian rhythms and is rhythmic under constant conditions suggesting they may be outputs of a circadian clock.

### Circadian expression of serpentine receptor 10

If malaria parasites are able to schedule the IDC they must respond to time-of-day information either through receptors or transporters. Serpentine receptor 10 (*sr10*: PF3D7_1215900), a member of a seven-transmembrane receptor family, was the top ranked receptor in the *P. falciparum* circadian gene list (ranked 21 out of all 495 genes sorted based on *q*-values) (**Fig. 3c**, **Supplementary Data 3**). Its orthologue in *P. chabaudi* (PCHAS_1433600) was also circadian in transcription in both matched and mismatched parasites (**Fig. 3c**, **Supplementary Data 1**). In *P. falciparum,* SR10 expression peaked at 8 h and 32 h post invasion, which corresponds to ring and late trophozoite stages of the IDC. In *P. chabaudi*, expression peaked at ZT14, corresponding to the late trophozoite stage.

Seven transmembrane domain-containing receptors/serpentine receptors/G protein coupled receptors (GPCRs) are the largest and most diverse group of membrane receptors and participate in a variety of physiological functions^41–44^. *P. falciparum* contains four serpentine receptor (SR) proteins; SR1, SR10, SR12 and SR25^45^. Of these, only *Pfsr10* showed circadian transcription while the rest showed periodicity closer to 48 h (**Supplementary Fig. 2a**). Expression profile of *Pfsr25* is not available in the microarray dataset from Bozdech et al (2003)^20^. However, microarray-based expression data (∼8h resolution) from another study^46^ revealed a single peak of expression during the entire IDC for *Pfsr25* with a periodicity closer to 48 h. Additionally, SR10 has been classified as a member of Class A serpentine receptors belonging to the hormonal receptor subclass based on the length of the N-terminal domain^45^ and classification by Inoue et al (2004)^47^. Of the four serpentine receptors, SR10 is also the only receptor that is present, not only in different *Plasmodium* spp., but also in other apicomplexans and distantly related organisms such as *Caenorhabditis elegans*, *Drosopihila melanogaster*, *Gallus gallus*, *Mus musculus*, *Homo sapiens* and *Arabidopsis thaliana* (data retrieved from OrthoMCL DB), although it has not been linked to circadian clocks in these organisms. The circadian transcription of *sr10* in both parasite species and in free-running conditions, coupled with its phylogenetic conservation across taxa, suggests the presence of a receptor-mediated signaling system in malaria parasites that receives time-of-day information from the host. We tested this hypothesis via a detailed analysis of SR10.

### Serpentine receptor 10 influences IDC duration

To investigate the functional role of SR10 *in vivo*, we disrupted the *sr10* gene in *P. chabaudi* by a double crossover homologous recombination strategy to generate *sr*10 deficient parasite clones (*sr10*KO) (**Supplementary Fig. 2b**). Functional disruption of *sr10* was verified by RNAseq analysis (**Supplementary Fig. 2c**). We then compared the IDC of wild type and two *sr10*KO clones by microscopic examination of thin blood smears (n= 4 per group), sampled every three hours over 48 h, starting from day 1 PI at 20.30 h (ZT13.5) (**Fig. 3d**). All infections of wild type and *sr10*KO clones (*sr10*KOA and B) were highly synchronous (amplitude ±(SE) for *P. chabaudi* wild type: 0.94±0.02, *P. chabaudi sr10*KOA: 0.79±0.02 and *sr10*KOB 0.93±0.03, **Fig. 3e**, **Supplementary Table 1**). Period estimates for the proportion of parasites at early trophozoite stage identified the IDC duration of both *sr10*KO clones to be ∼2h shorter (IDC duration 22.4 h) than the wild type (IDC duration 25.15; *p* < 0.0001, **Fig. 3f**).

Next, we investigated whether *sr10* also influences the IDC duration in another rodent malaria species i.e. *P. yoelii.* We generated an *sr10*KO clone in *P. yoelii* using a double crossover homologous recombination strategy (**Supplementary Fig. 2d**). As for *P. chabaudi*, we then compared wild type and *sr10*KO *P. yoelii* clones by microscopic examination of thin blood smears. The proportion of parasites at early trophozoite stage displayed weak circadian rhythmicity in both the wild type and *sr10*KO infections (Amplitude for *P. yoelii* wild-type: 0.30±0.02 and *P. yoelii sr10*KO: 0.31±0.01, **Fig. 3g**, **Supplementary Table 1**). As we find for the *P. chabaudi sr10*KO clones, knocking out *sr10* in *P. yoelii* shortens the duration of the IDC (*sr10*KO IDC duration 24.45 h; wild type IDC duration ∼28 h, **Fig. 3h**). Observing such similar biologically reproducible changes to IDC duration in two different experiments using two different malaria species strongly implicates *sr10* in the control of developmental progression through the IDC.

To explore how disruption of *sr10* affects the duration of the IDC we repeated the time-series RNAseq experiments on both wild type and *sr10*KO *P. chabaudi* parasites from 17 time points (however, data from the first three time points were excluded owing to a low number of mapped reads i.e. < 1 million paired ends mapped reads) sampled every three hours (n= 2 per time point) starting from day 2 PI (05.30 h, ZT22.5). Hierarchical clustering analysis identified biological replicates to be tightly clustered (**Supplementary Fig. 2e**). Transcripts with 24 h periodicity were identified following the approach deployed for the experiment using matched and mismatched *P. chabaudi* infections. A total of 3,620 and 2,886 genes showed ∼24 h rhythmicity in transcription in *P. chabaudi* wild type and *sr10*KO parasites respectively (*q* < 0.05; **Fig. 4a**, **Supplementary Data 4**). Principal component analysis of 14 time points identified that the first and third components of PCA (with a cyclic pattern in both the wild type and *sr10*KO parasites), accounted for > 85 % of total variance (**Supplementary Fig. 2f**). Comparison of wild type and *sr10*KO parasites revealed that 1,015 genes (19% of the total genes) lost transcriptional rhythmicity in *sr10*KO parasites (**Fig. 4b**, **Supplementary Data 4**). Further, 85% of the genes identified as being transcribed in a circadian manner in matched (wild type) *P. chabaudi* parasites from our first experiment were also circadian in the wild type parasites in this dataset (which are also matched). Whilst the additional rhythmically transcribed genes identified in wild type parasites in this dataset could be due to a longer time series, we found generally high concordance between the transcriptomes of infections initiated in the same way but in different laboratories. This lends support to the inference that the genes losing rhythmicity of transcription (henceforth called SR10-linked circadian genes or “SLCGs”) in *sr10*KO parasites is due to the loss of SR10.

**Fig. 4.**
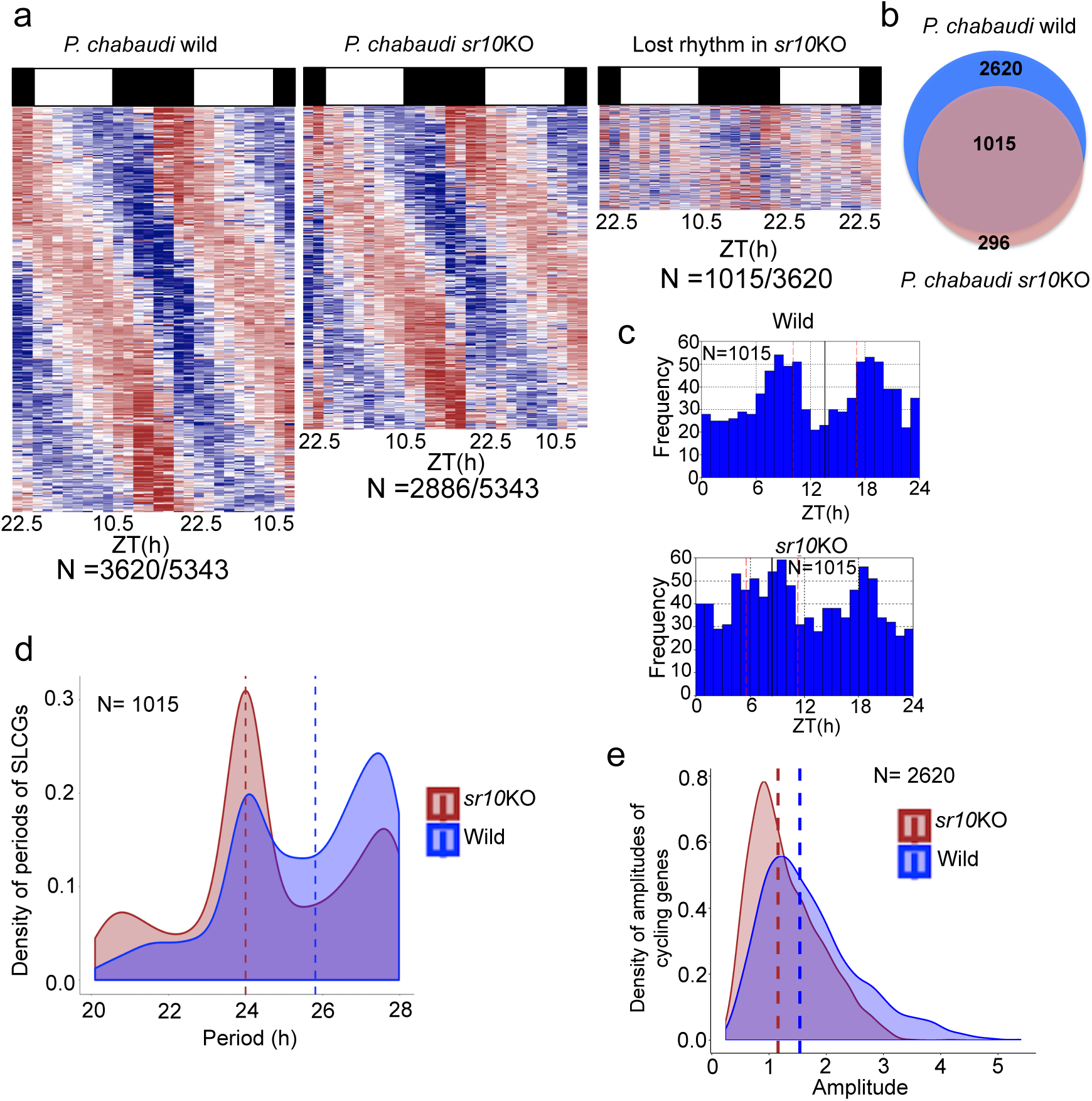
Disruption of SR10 affects the circadian transcriptome of *Plasmodium chabaudi*. **a)** Time series gene expression heatmap view of transcripts with 24 h rhythmicity in *P. chabaudi* wild type and *sr10*KO parasites. Right most heatmap shows the expression pattern of transcripts that lost rhythmicity in *sr*10KO parasites. Each row represents a single gene, sorted according to phase of maximum expression starting from first time point of sample collection. N represents number of circadian genes identified. **b)** Venn diagram of number of genes with 24 h rhythmic expression identified by both JTK and ARSER in wild type and *sr10*KO parasites. **c)** Phase distributions of genes with 24 h expression in wild type that lost 24 h rhythmicity in *sr10*KO parasites (SLCGs). The mean circular phase for each condition is indicated by a solid black line. N represents the number of cycling transcripts. Pink lines represent standard deviation of the mean circular phases. **d)** Transcripts that displayed with 24 h (putative “circadian”) rhythmicity only in wild parasites have median periods close to 26 h in wild parasites (blue dashed line) and 24 h in *sr10*KO parasites (brown dashed line). **e)** Genes that were rhythmic in both wild type and *sr10*KO parasites had a significantly lower mean amplitude in *sr10*KO parasites (1.15, brown dashed line) compared to wild parasites (1.53, *p* < 0.00001).

Examination of the SLCGs reveals a bimodal distribution pattern for peak transcription in wild-type parasites in which they are rhythmic (peaking at ZT 8 and ZT 19). This pattern was partially lost in *sr10*KO parasites in which the early peak displays a broader distribution (**Fig. 4c**). Further, the SLCGs exhibit a shorter periodicity in *sr10*KO (24 h) compared to wild type (25.81 h) parasites (**Fig. 4d**). Our intention is not to draw inference from the quantitative difference of periods (which is of limited utility for genes that lose rhythmicity), but to ascertain a qualitative comparison. However, the shorter periods in *sr10*KO parasites reflects the shorter periods observed in genes that lose rhythmicity in mismatched parasites. As genes that retained transcriptional rhythmicity in both matched and mismatched parasites, the genes that retained transcriptional rhythmicity in both wild type and *sr10*KO parasites (N=2,620) exhibited a significant reduction (*p* < 0.0001, unpaired student *t* test) of amplitude in *sr10*KO (1.15) compared to wild-type parasites (1.53) (**Fig. 4e**).

### SR10 regulates expression of multitude of pathways

Gene ontology (GO) analysis of the SLCGs showed enrichment for terms related to translation, RNA splicing, RNA and vesicle-mediated transport and purine and pyrimidine metabolism, indicating a broad effect of SR10 loss on parasite biology (**Supplementary Fig. 3**, **Supplementary Data 5**). Comparing differentially regulated genes for four different IDC stages (i.e. four time points: Day 2 ZT 16.5, Day 2 ZT 22.5, Day 3 ZT 4.5 and Day 3 ZT 10.5) in wild-type and *sr10*KO parasites (**Fig. 5a**, **Supplementary Data 6**) reveals that genes associated with: (i) protein translation are perturbed in ring stages; (ii) DNA replication and cell cycle associated processes are perturbed in early and late trophozoite stages; and (iii) microtubule based movement are perturbed in schizonts (**Fig. 5b**). Disruption of genes involved in DNA replication and cell cycle associated processes suggests SR10 influences IDC duration by altering the developmental rate of trophozoites. However, the perturbations in expression of rings and schizonts opens up the possibility that some, or all, of the IDC stages exhibit a shorter duration in *sr10*KO parasites.

**Fig. 5.**
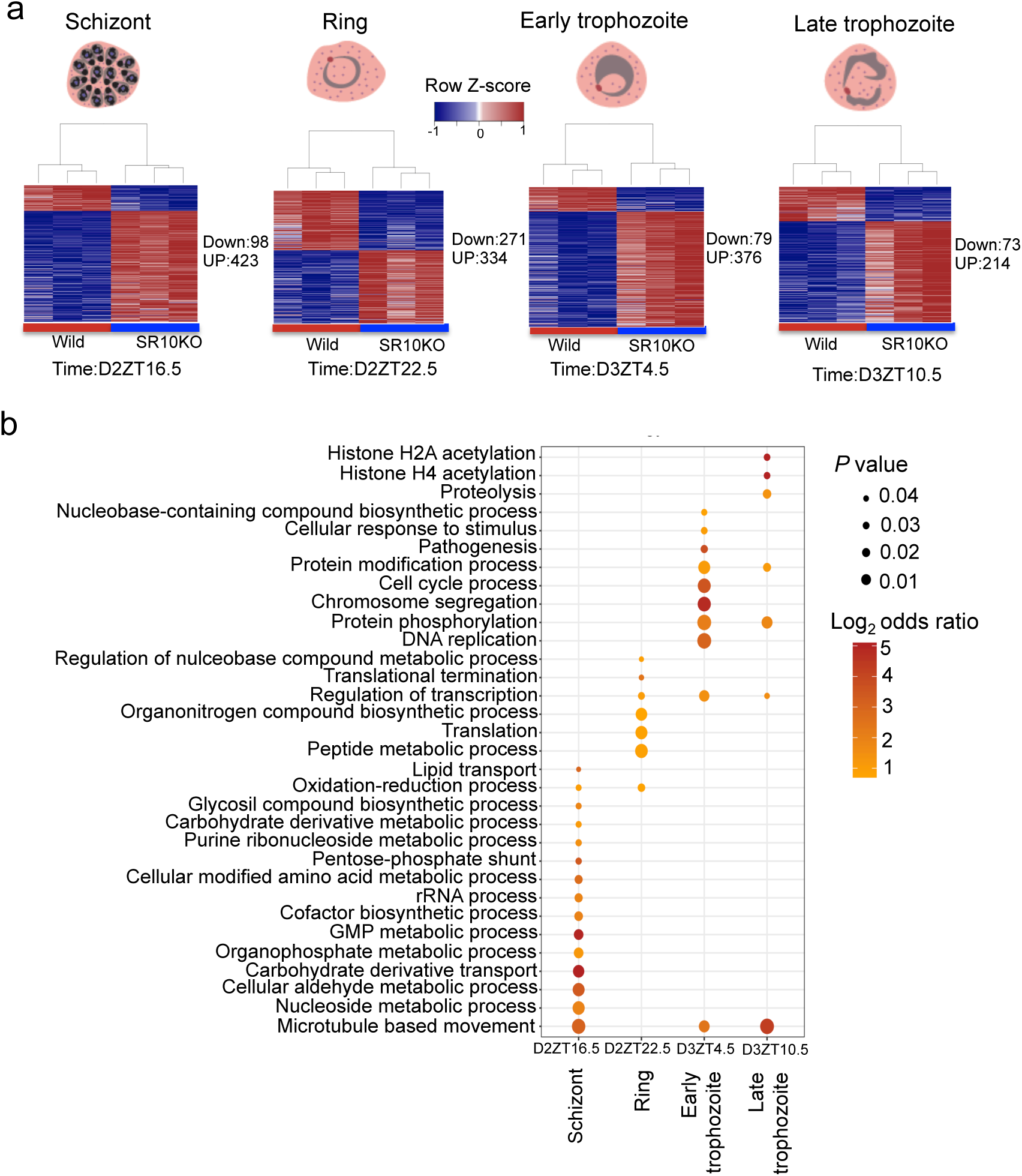
Knock out of *sr10* affects many biological processes. **a)** Differentially regulated genes were identified by comparing four matching time points of *sr10*KO and wild-type *Plasmodium chabaudi* parasites. Up and down represent differentially regulated genes with the false discovery rate corrected *p* < 0.05 and Log2 fold change < −1 for down-regulated genes and > 1 for up-regulated genes at each time point. The four time points analyzed represent four IDC stages as derived from examination of parasite morphology in thin blood smears. **b)** Gene ontology analysis of the differentially regulated genes within each time point. Manually curated functional annotations of biological processes (FDR <0.05) are represented and the colour spectrum represents the odds ratio.

We then compared the transcripts that lost rhythmicity in mismatched parasites (N = 1765) to the SLCGs (N = 1,015) to identify common genes. A total of 326 genes were shared (**Supplementary Fig. 4a**) suggesting that their transcription is shaped by both host rhythms and how the IDC is scheduled by the parasite. The shared genes were enriched for biological processes including energy metabolism, heme metabolic processes, and translation (**Supplementary Fig. 4b**).

### Disruption of *sr10* affects rhythmic expression of spliceosome machinery associated genes

The spliceosome is a large and dynamic ribonucleoprotein complex of five small nuclear ribonucleoproteins (snRNP) and over 150 multiple additional proteins that catalyze splicing of precursor mRNA in eukaryotes^48, 49^. Out of 85 genes (based on the Kyoto Encyclopedia of Genes and Genomes (KEGG) database mapping) expressing different spliceosomal proteins in *P. chabaudi*, 44 genes showed circadian transcription in wild type parasites, of which 26 are in the SLCG group (lost rhythmicity in *sr10*KO parasites) (**Fig. 6a**). They represent proteins of major spliceosome components including core spliceosomal protein members of snRNPs, prp19 complex and prp 19 related complexes.

**Fig. 6.**
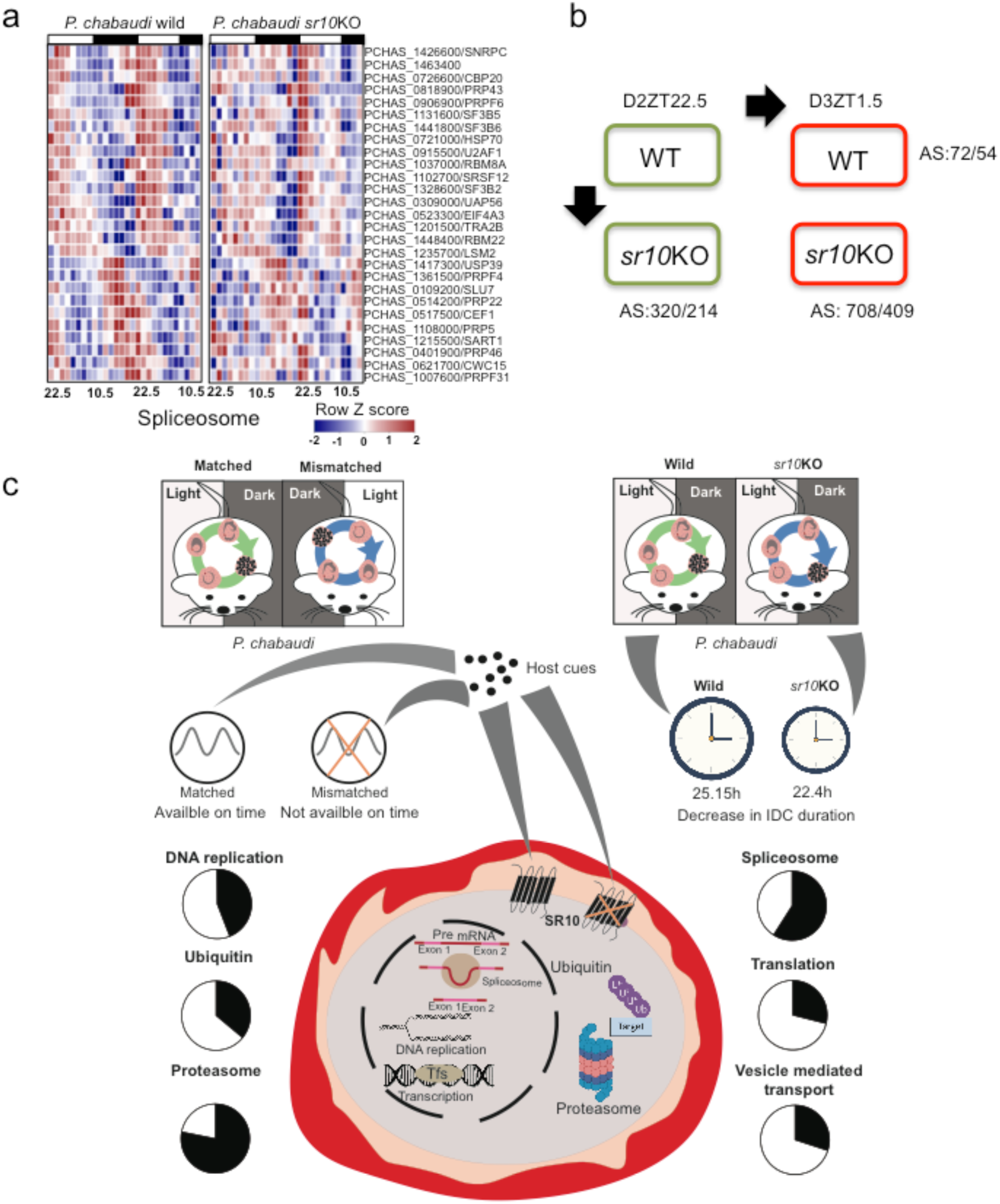
The cross-talks between the intraerythrocytic developmental cycle of *P. chabaudi* and the host rhythms. **a)** *sr10* knockout affects parasite spliceosome machinery. Heatmap illustrating the expression pattern of *sr10* knockout affected circadian genes involved in spliceosome pathway in *P. chabaudi* wild and *sr10*KO parasites. The list of genes was obtained by mapping the SLCGs to *P. chabaudi* spliceosome pathway represented in the KEGG database. Genes have been sorted based on phase of maximum expression. The colour scheme represents the row Z score. **b)** *sr10* knockout affects alternative splicing signature of the transcriptome. *sr10* knock out affects the alternative splicing signature of the parasite transcriptome. Two consecutive time points were compared between wild and *sr10*KO parasites to identify differential usage of exons. As a control two consecutive time points (Day 2 ZT 22.5 and Day 3 ZT1.5) from the same parasite strain were also compared. The number shown depict the number of differential exon usage events detected (*p* < 0.05). Two biological replicates per time point were used. Differential exon usage events were identified using DEXSeq^78^. **c)** Schematic figure summarizing the cross-talks between the intraerythrocytic developmental cycle schedule of *P. chabaudi* and the host rhythms. Parasite reschedules its IDC cycle when its developmental rhythms are mismatched with the host rhythms. Due to the mismatch, the parasites are devoid of host rhythmic cues on anticipated time. The parasites respond by losing rhythmic expression of genes associated with multiple biological processes as depicted in the pie-charts on the left. *P. chabaudi* serotonin receptor 10 (*sr10*) expresses rhythmically during the IDC and knocking out *sr10* in *P. chabaudi* reduces the IDC duration by ∼ 3h and also affects the rhythmic expression of genes associated with multiple biological processes as depicted in pie charts on the right. We speculate that SR10 may serve as one of the receptors through which the parasite receives the rhythmic cues of the host that controls the IDC duration of parasite. Black section within the pie-charts represent the percentage of rhythmic genes in each biological process that lost the rhythmic expression in the host-mismatched and *sr10*KO parasites.

Alternative splicing can regulate gene expression in signal dependent and tissue-specific manners^50^ and an emerging body of evidence links alternative splicing with the control of circadian regulatory networks in a variety of organisms, including *Drosophila melanogaster*^51^, *Neurospora crassa*^52–54^, *Arabidopsis*^55, 56^ and *Mus musculus*^57^. Alternative splicing has been reported to occur in Apicomplexans (including malaria parasites) for relatively few genes, covering only several percent of the total genes^58^.

Having observed that the loss of *sr10* modulates the transcription of genes associated with the spliceosome, we investigated whether it also affects the alternative-splicing (AS) signature using RNAseq analysis of AS in both wild type and *sr10*KO parasites. We considered two consecutive time-points, day 3 PI, time 05.30 GMT (ZT 22.5) and 08.30 (ZT 1.5) because they follow the time point when genes associated with spliceosome machinery are expressed maximally (ZT 16.5-19.5). Comparison of *sr10*KO and wild-type parasites identified 320 differential alternative splicing events covering 214 genes (*P*< 0.05) for ZT 22.5 and 708 differential alternative splicing events covering 409 genes (*p*< 0.05) for ZT1.5 (**Fig. 6b**, **Supplementary Data 7**). In a separate analysis, we compared two consecutive time points (ZT 22.5 and ZT 1.5) within each strain as controls, with an expectation of less differential alternative splicing events within compared to between wild-type and *sr10*KO parasites for the same time point. As expected, only 72 (54 genes) and 151 (114 genes) differential alternative splicing events were detected in each strain (**Figure 6B**, **Supplementary Data 7**). This suggests that our alternative splicing analysis was robust and that SR10 impacts the spliceosome machinery, resulting in differential alternative splicing patterns. GO-enrichment analysis of genes that showed differential alternative splicing events enriched with biological process terms such as translation, intracellular signal transduction and protein sumoylation (*Padj* < 0.05) of *sr10*KO compared to wild-type parasites. These observations collectively suggest that SR10 links host derived time-of-day information with how parasites schedule the IDC and regulate alternative splicing.

### Validation of circadian transcription pattern of genes through high-throughput real-time qPCR

We independently verified the transcription patterns of 87 genes that lost rhythmicity either in the mismatched parasites or in *sr10*KO parasites through high-throughput real-time qPCR using the BioMark^TM^ HD system (Fluidigm). A total of 58 genes, from the initial experiment comparing matched and mismatched *P. chabaudi* parasites, covering 11 randomly selected genes that represent multiple affected pathways were tested for their transcription in both groups (**Supplementary Fig. 4c**). Similarly, a total of 36 genes from the *sr10*KO experiment covering 12 randomly selected genes and representing multiple affected pathways were tested for their transcription in both *P. chabaudi* wild-type and *sr10*KO strains (**Supplementary Fig. 4d**). The gene datasets, generated by high-throughput real-time qRT-PCR using the BioMarkHD platform tightly correlated with the RNAseq transcription values (Spearman-rank correlation between 0.67 -0.95 for all the genes tested), thus independently validating our RNAseq analysis.

## DISCUSSION

How malaria parasites interact with host rhythms to establish and maintain rhythms during the IDC remain unknown. Our analyses, which were carried out using the rodent malaria parasites *P. chabaudi* and *P. yoelii in vivo,* and the human malaria parasite, *P. falciparum in vitro,* reveal an extensive transcriptome with 24 hour rhythmicity and suggest that coordination of the IDC with host rhythms is important for the parasites’ ability to undertake key cellular processes. This includes metabolic pathways, DNA replication, redox balance, the ubiquitin proteasome system, and alternative splicing (**Fig. 6c**). Rhythmicity in almost all of these processes persists in conditions in which parasites are not exposed to host rhythms, suggesting the presence of an endogenous time-keeping mechanism. We propose that, given its role in determining the duration of IDC and being a GPCR class of receptor, serpentine receptor 10 acts as a link between host circadian rhythms and parasite’s endogenous time-keeping / IDC scheduling mechanism.

Most genes (3,057 of 5,343) in the transcriptome of *P. chabaudi* in synchrony (matched) with the host circadian rhythm are transcribed with 24 hour periodicities, whilst this number drops to 1,824 when parasites are mismatched against the host circadian rhythm. Genes that lose rhythmicity are involved in diverse biological processes including glycolytic process, DNA replication, translation, ubiquitin/proteasome pathway and redox metabolism (**Supplementary Data 2**). This is not simply a consequence of mismatched parasites becoming desynchronised as they maintain synchrony during rescheduling (**Supplementary Fig. 1a**). Instead, disruption to these processes could be a result of stresses resulting from the IDC being misaligned to the host. If, for example, rescheduling parasites are unable to coincide the appearance of a particular IDC stage with a rhythmically provided resource it needs from the host, the parasite may be physiologically compromised and alternative pathways upregulated.

Regardless of the proximate cause, if such disruption impacts on the likelihood of completing the IDC or the fitness of merozoites produced by schizonts, it could explain the reduced replication rate previously observed in mismatched *P. chabaudi*^15, 16^. Whilst caution needs to be employed in interpreting the periodicities of genes that lost rhythmicity of transcription, on the whole, they were approximately 1 hour shorter in mismatched than matched parasites. This difference is supported by the observation that mismatched parasites were rescheduling by on average approximately 1.5 hour every IDC. Why some genes retain rhythmicity (1,292) and others don’t (1,765) is unclear. Further, why 532 genes – all associated with RNA processing/splicing - became rhythmic as a consequence of mismatch is unclear but raises the possibility of an alternative set of IDC and/or timekeeping genes.

The IDC of *P. chabaudi* is approximately 24 hours in duration, making it difficult to distinguish genes associated simply with particular IDC stages from genes associated with an endogenous time-keeping mechanism or its outputs. Thus, *P. falciparum* cultured under constant conditions was used to separate IDC from putative clock/clock controlled genes. We found that 495 *P. falciparum* genes exhibit ∼ 24 hour transcriptional rhythms and many of these genes are associated with metabolic processes (carbohydrate metabolism, DNA replication and the ubiquitin proteasome system) affected in *P. chabaudi* by mismatch with host rhythm.

The presence of “free-running” rhythms is consistent with an endogenous oscillator. That the transcription of these genes becomes disrupted by mismatch is analogous to “jet lag”; the temporal misalignment of processes in different organs/tissues whilst a clock regains coordination (entrains) to a phase-shift in its time-cue (Zeitgeber). Such misalignment can manifest as a dampening (reduction in amplitude) of rhythmicity of transcription of genes driven by the clock^22, 59^. In concordance with this, we observed a reduction in the mean amplitude of transcripts of genes that remain rhythmic in matched, mismatched (**Fig. 1d**), and in wild type and *sr10*KO parasites (**Fig. 4e**).

Observing that the ubiquitin-proteasome system is disrupted in mismatched parasites is also consistent with parasites possessing an endogenous oscillator as the ubiquitin-proteasome system plays a role in regulating clock components and their outputs in many taxa^28^. The timing of transcription of ubiquitin-proteasome system genes in mismatched *P. chabaudi* corresponds to the final stage of the IDC, and so could be disrupted as a mechanism to facilitate a shorter IDC or as consequence of the wrong IDC stage receiving time-of-day information from the host, or both. Similarly, if the signal to begin DNA replication, acquire glucose, and / or manage redox states, is host derived, mismatched parasites will be at the wrong IDC stage to respond appropriately.

The strongest suggestion that the parasite is capable of scheduling the IDC comes from observations that SR10; (i) determines IDC duration in *P. chabaudi* and *P. yoelii* (**Fig. 6c**); (ii) has a 24 h transcription rhythm in *P. falciparum* and in both matched and mismatched *P. chabaudi;* (iii) peaks at trophozoite stage in both *P. falciparum* and *P.chabaudi;* (iv) regulates rhythmicity in gene transcription for several of the processes whose genes lost transcription rhythmicity in mismatched *P. chabaudi* (**Fig. 6c**); and (v) determines the period of transcripts for genes that lose rhythmicity when it is disrupted.

We also observed rhythmic transcription of genes associated with histone modification and the control of transcription and translation (**Figs 2a, 3a, 5b** and **Supplementary** **fig. 3**). These processes are considered central to the circadian organization of the transcriptome^60–62^. Thus, we propose that SR10 acts as a link between time-of-day information provided by host rhythms and the parasites’ endogenous time-keeping mechanism that schedules the IDC.

In summary, we reveal that coordination with host circadian rhythms is central to the schedule of transcription of genes associated with diverse processes underpinning IDC progression, and ultimately, replication. Our data are consistent with some of the criteria required to demonstrate an endogenous time-keeping ability, suggesting that malaria parasites are at least in part responsible for scheduling their IDC. Further work should examine whether other features of an endogenous clock exist and identify the interaction partners of SR10 that link it with host time-of-day.

Taking all of our observations together, we propose that: (i) malaria parasites benefit from coordination with the hosts circadian rhythm, (ii) malaria parasites are capable of sensing the timing of their host’s feeding rhythm and adjust the speed of the IDC, possibly by altering the duration of the trophozoite stage, to facilitate coordination with nutrient supplies. The IDC underpins malaria parasites’ capacity to undergo rapid asexual replication and cause severe disease, and fuels transmission of the disease. Thus, uncovering the components of the parasites’ time-keeping mechanisms and the signaling system that links it to IDC progression may uncover novel intervention strategies.

## METHODS

### Ethics statement

All animal procedures for the parasite mismatching study were performed in accordance with the UK Home office regulations (Animals Scientific Procedures ACT 1986; project license number 70/8546) and approved by the University of Edinburgh. All the procedures for the *sr10*KO study were performed in strict accordance with the Japanese Humane Treatment and Management of Animal Law (Law No. 105 dated 19 October 1973 modified on 2 June 2006), and the Regulation on Animal Experimentation at Nagasaki University, Japan. The protocol was approved by the Institutional Animal Research Committee of Nagasaki University (permit: 12072610052). The animal experiments conducted in Edinburgh and Nagasaki were officially exempted from additional IBEC clearances in KAUST. All the procedures to perform work on different parasite materials used in this study was approved by IBEC in KAUST (IBEC number: 19IBEC12)

### Parasites and hosts: Experimental design and data collection for host rhythm mismatching experiments

Hosts were 6-8 weeks old female MF1 mice housed in groups of five at 21°C with food and drinking water supplemented with 0.05 % para-aminobenzoic acid (PABA, to supplement parasite growth) provided *ad libitum*. All experimental infections were initiated by intravenous injection of 1 × 10^7^ *P. chabaudi chabaudi* clone AS^63^ parasitized red blood cells (per ml). Parasites at ring stage were collected at ZT0.5 (08:00 GMT) from donor mice housed in standard LD light conditions (Lights ON 07:30; Lights OFF 19:30 GMT) and immediately used to infect two groups of experimental mice. One group (termed “matched”) were entrained to the same photoperiod as the donor mice, generating infections in which parasite and host rhythms were in the same phase. The other group (termed “mismatched”) were entrained to reverse light (Lights OFF 07:30 and Lights ON 19:30 GMT) resulting in infections in which the parasite is out of phase with the host by 12 hours.

Samples were collected every three hours, over a period of 30 hours, from 09:00 GMT on day 4 to 15:00 GMT on day 5 post infection. This corresponds to a starting time of ZT1.5 for matched, and ZT13.5 for mismatched infections. At each sampling point, four mice from each group (matched/mismatched) were sacrificed and thin blood smears and RBC counts (*via* flow cytometry; Beckman Coulter) were taken by tail bleeds, and 50µl of blood was taken *via* cardiac puncture (added to 200µl of RNAlater and frozen at −80C) for RNAseq analysis. The developmental rhythm of parasites was assessed from blood smears, in which the number of parasites at each of three morphologically distinct stages (ring stage trophozoite (hereby referred to as ‘ring stage’), early (small) trophozoite stage, late (large) trophozoites and schizont; differentiated based on parasite size, the size and number of nuclei and the appearance of haemozoin) was recorded, as described before^13^.

### Experimental design and data collection for sr10 knockout experiment

The *sr10* gene knockout and subsequent experiments were performed with the *P. chabaudi* AS clone. Routine maintenance of the parasites was performed in ICR female mice (6-8 weeks old) and rhythmicity was assessed in groups of female CBA inbred mice (6-8 weeks old). Mice (SLC Inc., Shizuoka, Japan) were housed at 23°C with 12hr light-dark cycle (lights-off: 19:00 h and lights-on: 07:00 h) and fed on a maintenance diet with 0.05% PABA-supplemented water.

Ring stage *sr10*KO (clone A and B) and wild type *P. chabaudi* parasites were sub-inoculated into groups of four CBA mice each (1 x 10^6^ parasitized RBCs per mouse) by intravenous injection at 20:30 hrs corresponding to ZT13.5, day 0. Starting at ZT13.5 on day 1 post-infection, blood smears were taken for both groups every three hours for 48 hrs producing a total of 17 time points. Blood smears were briefly fixed with 100% methanol and stained with Giemsa’s solution. The parasitic patterns were then recorded based on classification of the parasitic forms into four stages-rings (ring stage trophozoite), early (small) trophozoites, late (large) trophozoites and schizonts. Same procedures were adopted for *P. yoelii* wild type (17×1.1pp) and *sr10*KO clones to obtain time series phenotype data using blood smears. Blood microsamples were also collected at these timepoints for time-series gene expression analysis^64^. Briefly, 20µl of blood was collected *via* tail snip at each time point, washed in PBS and immediately treated with 500 uL TRIzol reagent and stored shortly at 4°C and for long term at −80°C.

### Plasmid construction to modify the sr10 gene locus in Plasmodium chabaudi and Plasmodium yoelii

Plasmids were constructed using the MultiSite Gateway cloning system (Invitrogen). For *P. chabaudi*, One thousand base pair long regions at the 5′ and 3′ UTRs of *Pchsr*10 were PCR-amplified from *P. chabaudi* with *att*B-flanked primers, *Pchsr10*-5U.B1.F and *Pch*-5U.B2.R to yield *att*B1-*Pchsr10*-5U-*att*B2 fragment, and *Pchsr10*-3U.B4.F and *Pchsr10*-3U.B1r.R to yield *att*B4-*Pchsr10*-3U-*att*B1r fragment. The *att*B1-*Pchsr10*-5U-*att*B2 and *att*B4-*Pchsr10*-3U-*att*B1r products were then subjected to independent BP recombination reactions with pDONR221 (Invitrogen) and pDONRP4P1R (Invitrogen) to generate pENT12-*Pchsr10*-5U and pENT41-*Pchsr10*-3U entry clones, respectively. All BP reactions were performed using the BP Clonase II enzyme mix (Invitrogen) according to the manufacturer’s instructions. pENT12-*Pchsr10*-5U, pENT41-*Pchsr10*-3U and linker pENT23-3Ty entry plasmids were subjected to LR recombination reaction (Invitrogen) with a destination vector pDST43-HDEF-F3 (that contains the pyrimethamine resistant gene selection cassette *h*DHFR) to yield knockout construct pKO-*Pchsr10*. LR reactions were performed using the LR Clonase II Plus enzyme mix (Invitrogen) according to the manufacturer’s instructions. *P. yoelii* 17x1.1pp *sr10* knockout parasites were also generated using the same procedures adopted for *P. chabaudi* using *Pysr10*-5U.B1.F and *Pysr10*-5U.B2.R to yield *att*B1-*Pysr10*-5U-*att*B2 fragment, and *Pysr10*-3U.B4.F PySR10-3U.B1r.R to yield *att*B4-*Pysr10*-3U-*att*B1r fragment from *P. yoelii* 17×1.1pp gDNA. These fragments were then used in independent BP reactions and subsequent LR reactions as above to generate pKO-*Pysr10* construct. All primers used are listed in **Supplementary Data 8**.

### Parasite transfection

Schizonts from *P. yoelii* and *P. chabaudi*-infected mice were enriched by centrifugation over a Histodenz density cushion. Histodenz™ (Sigma-Aldrich, St. Louis, MO) solution was prepared as 27.6 g/100 mL in Tris-buffered solution (5 mM Tris-HCl, 3 mM KCl, and 0.3 mM CaNa2-EDTA, pH 7.5) and then diluted with equal volume of RPMI1640-based incomplete medium containing 25 mM HEPES and 100 mg/L of hypoxanthine^65^. The schizont-enriched parasites were transfected using a Nucleofector™ 2b device (Lonza Japan) as described^66^ using 20 μg of linearized plasmids for each transfection. Stable transfectants were selected by oral administration of pyrimethamine (0.07mg/mL) and cloned by limiting dilution in mice. Stable integration of plasmids in the parasite genome was confirmed by PCR and sequencing.

#### Time series gene expression analysis of parasites using RNAseq

Total RNA was isolated from TRIzol treated samples according to the manufacturer’s instructions (Life Technologies). Strand-specific mRNA libraries were prepared from total RNA using TruSeq Stranded mRNA Sample Prep Kit LS (Illumina) according to the manufacturer’s instructions. Briefly, at least 100ng of total RNA was used as starting material to prepare the libraries. PolyA+ mRNA molecules were purified from total RNA using oligo-T attached magnetic beads. First strand synthesis was performed using random primers followed by second strand synthesis where dUTP were incorporated in place of dTTP to achieve strand-specificity. Ends of the double stranded cDNA molecules were adaptor ligated and the libraries amplified by PCR for 15 cycles. Libraries were sequenced on Illumina HiSeq 4000 platform with paired-end 100/150bp read chemistry according to manufacturer’s instructions.

#### Real-time quantitative reverse transcriptase PCR analysis

Total RNA was treated with TURBO Dnase according to the manufacturer’s instructions (Thermo Fischer Scientific) to eliminate DNA contamination. The absence of DNA in RNA samples was confirmed by inability to detect DNA after 40 cycles of PCR with HSP40, family A (PCHAS_0612600) gene primers in a 7900HT fast real-time PCR system (Applied Biosystems) with the following cycling conditions: 95**°**C for 30 sec followed by 40 cycles of 95**°**C 2 sec; 60**°**C for 25 sec. *P. chabaudi* DNA was used as positive control. For both the mismatching and SR10 experiments, gene expression profiles were obtained for a total of 87 genes from eight time-points. Two biological replicates per experimental condition and two technical replicates per biological replicate were run on a Biomark HD microfluidic quantitative RT-PCR platform (Fluidigm) to measure the expression level of genes. For the SR10 experiment, first strand cDNA synthesis was performed using reverse transcription master mix according to the manufacturer’s instructions (Fluidigm) and for the mismatching experiment, first strand cDNA synthesis was performed using a High-Capacity cDNA reverse transcription kit according to the manufacturer’s instructions (Thermo Fisher Scientific). Pre-amplification of target cDNA was performed using a multiplexed, target-specific amplification protocol (95**°**C for 15 sec, 60**°**C for 4 min for a total of 14 cycles). The pre-amplification step uses a cocktail of forward and reverse primers of targets (genes of interest) under study to increase the number of copies to a detectable level. Products were diluted 5-fold prior to amplification using SsoFast EvaGreen Supermix with low ROX and target specific primers in 96.96 Dynamic arrays on a Biomark HD microfluidic quantitative RT-PCR system (Fluidigm). Expression data for each gene were retrieved in the form of Ct values. Normalization of transcript expression level was carried out using *P. chabaudi* HSP40, family A (PCHAS_0612600) gene that was found to be non-circadian in expression in all the strains used. All primers used are listed in **Supplementary Data 8**.

#### Analysis of parasite developmental rhythmicity and periodicity

The early trophozoite stage was used as the marker stage because this stage is both easily identifiable and has developmental duration similar to other stages. Rhythmicities in the proportion of parasites at early trophozoite stages was determined by Fourier transformed harmonic regression in Circwave^67^. A cosine wave was fitted to data from each individual infection and compared to a straight line at the mean *via* an F-test. The period was allowed to change between 18 and 30 hours (i.e. 24hr ± two sampling periods for the host rhythm mismatch experiment and one sampling period for the analysis of *sr10*KO strains) and the fit was considered significant if the adjusted (to account for multiple tests for different periods) *p* value was greater than the alpha of 0.017. General linear models were used to determine if the characteristics of rhythms varied according to treatment and parasite strain (using the package lme4 in R version 3.4.0).

### Transcriptome sequencing and analysis

RNASeq read quality was assessed using FASTQC quality control tool (http://www.bioinformatics.babraham.ac.uk/projects/fastqc). Read trimming tool Trimmomatic^68^ was used to remove low quality reads and Illumina adaptor sequences. Reads smaller than 36 nucleotides long were discarded. Quality trimmed reads were mapped to *P. chabaudi chabaudi* AS reference genome (release 28 in PlasmoDB-http://www.plasmoddb.org) using TopHat2 (version 2.0.13)^69^ with parameters “ –no-novel-juncs –library-type fr-firststrand”. Gene expression estimates were made as raw read counts using the Python script ‘HTSeq-count’ (model type – union, http://www-huber.embl.de/users/anders/HTSeq/)^70^. Count data were converted to counts per million (cpm) and genes were filtered if they failed to achieve a cpm value of 1 in at least 30% of libraries per condition. Library sizes were scale-normalized by the TMM method using EdgeR software^71^ and further subjected to linear model analysis using the voom function in the limma package^72^. Differential expression analysis was performed using DeSeq2^73^. Genes with fold change greater than two and false discovery rate corrected p-value (Benjamini-Hochberg procedure) < 0.05 were considered to be differentially expressed.

### Identification of circadian transcripts in *Plasmodium chabaudi*

Circadian transcripts were identified using two programs, JTK-Cycle^74^ and ARSER^75^ implemented in MetaCycle^76^, an integrated R package with parameters set to fit time-series data to exactly 24-h periodic waveforms. While JTK Cycle uses a non-parametric test called the Jonckheere-Terpstra test to detect rhythmic transcripts^74^, ARSER uses “autoregressive spectral estimation to predict an expression profile’s periodicity from the frequency spectrum and then models the rhythmic patterns by using a harmonic regression model to fit the time-series” ^75^. For both the programs voom-TMM normalized count data was used as input data. A gene was considered cyclic if both the programs identified it as a circadian transcript with significance bounded by *p* < 0.05 for the parasite and host rhythms mismatching experiment where 11 time points separated by 3h were used and by q < 0.05 for the SR10 experiment where 14 time points separated by 3h were used. We used the data from only 14 out of 17 time points. The first three time-points having been excluded owing to low numbers of mapped reads to *P. chabaudi* (< 1 million). The output from ARSER concerning amplitude, phase and period of circadian transcripts was used for further analysis.

Time points and biological replicates were clustered using hierarchical clustering in R software environment with Pearson correlations from normalized count values as input and ‘ward.D2’ agglomerative hierarchical clustering procedure was used for cluster generation. The circular and linear histogram plots representing phase distribution of cycling transcripts were generated using Oriana (www.kovcomp.co.uk/oriana/). PCA was performed on voom-TMM normalized data using *princomp* in R software environment.

To determine the FDR of circadian transcripts in the host-circadian rhythm mismatching experiment, the time points of collection were randomly permuted 1000 times and the number of circadian transcripts was assessed for each of the permutations by both the programs used. A similar approach to determine the FDR was used by^10^. This was done for both matched and mismatched parasite datasets.

### Identification and analysis of circadian transcripts in *Plasmodium falciparum*

Microarray based stage-specific expression data covering *P. falciparum* intra-erythrocytic developmental stages sampled at 1 hour resolution for 48 hours was obtained from a previous study^20^. The data had two odd time points (TP) missing (TP 23 and TP 29) so only even time points were considered. The final data had 24 time points with a 2 h sampling rate. Circadian transcripts were identified following the same protocol as for *P. chabaudi*. *P. chabaudi* and *P. falciparum* gene ontology terms were downloaded from the UniProt gene ontology annotation database (https://www.ebi.ac.uk/GOA). Circadian genes were segregated into 12 groups based on their phase of maximum expression as determined from ARSER output and gene ontology enrichment analysis was performed on each groups using GOstats R package^77^. In the case of *P. falciparum*, GO-enrichment analysis was performed on all the identified circadian transcripts. GO terms were considered only if statistical tests showed FDR corrected *p* < 0.05. Odds ratio was calculated by dividing the occurrence for GO term in the input list to the occurrence for GO term in the reference set (i.e. whole genome).

### Identification of differential alternative splicing events

Differential alternative splicing events in terms of differential exon usage were detected using the DEXSeq program v 1.20.02^78^ with modified scripts as reported in Yeoh et al. (2015)^79^. The *p* value significance level was set to 0.05 for the identification of differential exon usage. Comparison was made between *P. chabaudi* wild and the SR10 knocked out strains for two time points *i.e.* Day 3, ZT 21 and Day 3, ZT 0/24 post-infection. For these two time points, RNASeq read depth was increased by performing additional rounds of sequencing in order to detect AS events more reliably. Two biological replicates per time-point were used.

## DATA AND SOFTWARE AVAILABILITY

RNASeq data sets associated with this study have been submitted to the gene expression omnibus under accession number GSE132647. See also Supplementary Table 1 and 3.

## Supporting information

Supplemental Filles

## ACKNOWLEDGEMENTS

The project was supported by a faculty baseline funding (BAS/1/1020-01-01) from the King Abdullah University of Science and Technology (KAUST) to AP. RC is supported by Japanese Society for the Promotion of Science (JSPS), Japan Grant-in-Aid for Scientific Research Nos. 24255009, 25870525, 16K21233 and 19K07526. SER and AJOD are supported by Wellcome (202769/Z/16/Z; 204511/Z/16/Z), the Royal Society (UF110155; NF140517) and the Human Frontier Science Program (RGP0046/2013). The authors thank the staff of the Bioscience Core Laboratory in KAUST for sequencing RNAseq libraries and all members of the Reece lab at the University of Edinburgh and pathogen genomics lab at KAUST for assistance during the experiments. This work was partly conducted at the Joint Usage / Research Center for Tropical Disease, Institute of Tropical Medicine, Nagasaki University, Japan.

## AUTHOR CONTRIBUTIONS

Conceptualization, A.P. and S.E.R.; Methodology, A.P., S.E.R., A.K.S., A.J.O.D., H.M.A., A.R., R.C. and O.K.; Investigation, A.K.S., A.R., A.J.O.D., R.C., and H.M.A.; Formal analysis, A.K.S., A.R., A.J.O.D., H.M.A., A.K., A.M.A.H., F.B.R. and H.R.A.; Writing-Original Draft, A.K.S., S.E.R and A.P.; Writing – Review & Editing, A.K.S., S.E.R, R.C. and A.P.; Funding Acquisition, S.E.R. and A.P.; Resources, S.E.R., R.C. and A.P.; Supervision, A.P.

## REFERENCES

1. Sharma, V. K. Adaptive significance of circadian clocks. Chronobiol Int 20, 901–919 (2003).

2. Bell-Pedersen, D. et al. Circadian rhythms from multiple oscillators: lessons from diverse organisms. Nat Rev Genet 6, 544–556, doi:10.1038/nrg1633 (2005).

3. Edgar, R. S. et al. Peroxiredoxins are conserved markers of circadian rhythms. Nature 485, 459–464, doi:10.1038/nature11088 (2012).

4. Rosbash, M. The implications of multiple circadian clock origins. PLoS Biol 7, e62, doi:10.1371/journal.pbio.1000062 (2009).

5. Young, M. W. & Kay, S. A. Time zones: a comparative genetics of circadian clocks. Nat Rev Genet 2, 702–715, doi:10.1038/35088576 (2001).

6. O’Neill, J. S. et al. Circadian rhythms persist without transcription in a eukaryote. Nature 469, 554–558, doi:10.1038/nature09654 (2011).

7. O’Neill, J. S. & Reddy, A. B. Circadian clocks in human red blood cells. Nature 469, 498–503, doi:10.1038/nature09702 (2011).

8. Reece, S. E., Prior, K. F. & Mideo, N. The Life and Times of Parasites: Rhythms in Strategies for Within-host Survival and Between-host Transmission. J Biol Rhythms 32, 516–533, doi:10.1177/0748730417718904 (2017).

9. Westwood, M. L. et al. The evolutionary ecology of circadian rhythms in infection. Nat Ecol Evol 3, 552–560, doi:10.1038/s41559-019-0831-4 (2019).

10. Rijo-Ferreira, F., Pinto-Neves, D., Barbosa-Morais, N. L., Takahashi, J. S. & Figueiredo, L. M. Trypanosoma brucei metabolism is under circadian control. Nat Microbiol 2, 17032, doi:10.1038/nmicrobiol.2017.32 (2017).

11. Hawking, F. The clock of the malaria parasite. Sci Am 222, 123–131 (1970).

12. Hirako, I. C. et al. Daily Rhythms of TNFalpha Expression and Food Intake Regulate Synchrony of Plasmodium Stages with the Host Circadian Cycle. Cell Host Microbe, doi:10.1016/j.chom.2018.04.016 (2018).

13. Prior, K. F. et al. Timing of host feeding drives rhythms in parasite replication. PLoS Pathog 14, e1006900, doi:10.1371/journal.ppat.1006900 (2018).

14. Reece, S. E. & Prior, K. F. Malaria Makes the Most of Mealtimes. Cell Host Microbe 23, 695–697, doi:10.1016/j.chom.2018.05.015 (2018).

15. O’Donnell, A. J., Schneider, P., McWatters, H. G. & Reece, S. E. Fitness costs of disrupting circadian rhythms in malaria parasites. Proc Biol Sci 278, 2429–2436, doi:10.1098/rspb.2010.2457 (2011).

16. O’Donnell, A. J., Mideo, N. & Reece, S. E. Disrupting rhythms in Plasmodium chabaudi: costs accrue quickly and independently of how infections are initiated. Malar J 12, 372, doi:10.1186/1475-2875-12-372 (2013).

17. Hott, A. et al. Artemisinin-resistant Plasmodium falciparum parasites exhibit altered patterns of development in infected erythrocytes. Antimicrob Agents Chemother 59, 3156–3167, doi:10.1128/AAC.00197-15 (2015).

18. Mok, S. et al. Artemisinin resistance in Plasmodium falciparum is associated with an altered temporal pattern of transcription. BMC Genomics 12, 391, doi:10.1186/1471-2164-12-391 (2011).

19. Mok, S. et al. Drug resistance. Population transcriptomics of human malaria parasites reveals the mechanism of artemisinin resistance. Science 347, 431–435, doi:10.1126/science.1260403 (2015).

20. Bozdech, Z. et al. The transcriptome of the intraerythrocytic developmental cycle of Plasmodium falciparum. PLoS Biol 1, E5, doi:10.1371/journal.pbio.0000005 (2003).

21. Hoo, R. et al. Integrated analysis of the Plasmodium species transcriptome. EBioMedicine 7, 255–266, doi:10.1016/j.ebiom.2016.04.011 (2016).

22. Archer, S. N. et al. Mistimed sleep disrupts circadian regulation of the human transcriptome. Proc Natl Acad Sci U S A 111, E682–691, doi:10.1073/pnas.1316335111 (2014).

23. Chen, C. Y. et al. Effects of aging on circadian patterns of gene expression in the human prefrontal cortex. Proc Natl Acad Sci U S A 113, 206–211, doi:10.1073/pnas.1508249112 (2016).

24. Bailey, S. M., Udoh, U. S. & Young, M. E. Circadian regulation of metabolism. J Endocrinol 222, R75–96, doi:10.1530/JOE-14-0200 (2014).

25. Milev, N. B. & Reddy, A. B. Circadian redox oscillations and metabolism. Trends Endocrinol Metab 26, 430–437, doi:10.1016/j.tem.2015.05.012 (2015).

26. Kalsbeek, A., la Fleur, S. & Fliers, E. Circadian control of glucose metabolism. Mol Metab 3, 372–383, doi:10.1016/j.molmet.2014.03.002 (2014).

27. Mancio-Silva, L. et al. Nutrient sensing modulates malaria parasite virulence. Nature 547, 213–216, doi:10.1038/nature23009 (2017).

28. Stojkovic, K., Wing, S. S. & Cermakian, N. A central role for ubiquitination within a circadian clock protein modification code. Front Mol Neurosci 7, 69, doi:10.3389/fnmol.2014.00069 (2014).

29. Kamura, T. et al. Rbx1, a component of the VHL tumor suppressor complex and SCF ubiquitin ligase. Science 284, 657–661 (1999).

30. Ohta, T., Michel, J. J., Schottelius, A. J. & Xiong, Y. ROC1, a homolog of APC11, represents a family of cullin partners with an associated ubiquitin ligase activity. Mol Cell 3, 535–541 (1999).

31. Seol, J. H. et al. Cdc53/cullin and the essential Hrt1 RING-H2 subunit of SCF define a ubiquitin ligase module that activates the E2 enzyme Cdc34. Genes Dev 13, 1614–1626 (1999).

32. Skowyra, D. et al. Reconstitution of G1 cyclin ubiquitination with complexes containing SCFGrr1 and Rbx1. Science 284, 662–665 (1999).

33. Dong, G. et al. Elevated ATPase activity of KaiC applies a circadian checkpoint on cell division in Synechococcus elongatus. Cell 140, 529–539, doi:10.1016/j.cell.2009.12.042 (2010).

34. Feillet, C. et al. Phase locking and multiple oscillating attractors for the coupled mammalian clock and cell cycle. Proc Natl Acad Sci U S A 111, 9828–9833, doi:10.1073/pnas.1320474111 (2014).

35. Kowalska, E. et al. NONO couples the circadian clock to the cell cycle. Proc Natl Acad Sci U S A 110, 1592–1599, doi:10.1073/pnas.1213317110 (2013).

36. Laranjeiro, R. et al. Cyclin-dependent kinase inhibitor p20 controls circadian cell-cycle timing. Proc Natl Acad Sci U S A 110, 6835–6840, doi:10.1073/pnas.1217912110 (2013).

37. Masri, S., Cervantes, M. & Sassone-Corsi, P. The circadian clock and cell cycle: interconnected biological circuits. Curr Opin Cell Biol 25, 730–734, doi:10.1016/j.ceb.2013.07.013 (2013).

38. Reddy, A. B. & Rey, G. Metabolic and nontranscriptional circadian clocks: eukaryotes. Annu Rev Biochem 83, 165–189, doi:10.1146/annurev-biochem-060713-035623 (2014).

39. Sargeant, T. J. et al. Lineage-specific expansion of proteins exported to erythrocytes in malaria parasites. Genome Biol 7, R12, doi:10.1186/gb-2006-7-2-r12 (2006).

40. Otto, T. D. et al. New insights into the blood-stage transcriptome of Plasmodium falciparum using RNA-Seq. Mol Microbiol 76, 12–24, doi:10.1111/j.1365-2958.2009.07026.x (2010).

41. Buck, L. & Axel, R. A novel multigene family may encode odorant receptors: a molecular basis for odor recognition. Cell 65, 175–187 (1991).

42. Firestein, S. The good taste of genomics. Nature 404, 552–553, doi:10.1038/35007167 (2000).

43. Howard, A. D. et al. Orphan G-protein-coupled receptors and natural ligand discovery. Trends Pharmacol Sci 22, 132–140 (2001).

44. Lee, D. K., George, S. R., Evans, J. F., Lynch, K. R. & O’Dowd, B. F. Orphan G protein-coupled receptors in the CNS. Curr Opin Pharmacol 1, 31–39 (2001).

45. Madeira, L. et al. Genome-wide detection of serpentine receptor-like proteins in malaria parasites. PLoS One 3, e1889, doi:10.1371/journal.pone.0001889 (2008).

46. Rovira-Graells, N. et al. Transcriptional variation in the malaria parasite Plasmodium falciparum. Genome Res 22, 925–938, doi:10.1101/gr.129692.111 (2012).

47. Inoue, Y., Ikeda, M. & Shimizu, T. Proteome-wide classification and identification of mammalian-type GPCRs by binary topology pattern. Comput Biol Chem 28, 39–49 (2004).

48. Jurica, M. S. & Moore, M. J. Pre-mRNA splicing: awash in a sea of proteins. Mol Cell 12, 5–14 (2003).

49. Will, C. L. & Luhrmann, R. Spliceosome structure and function. Cold Spring Harb Perspect Biol 3, doi:10.1101/cshperspect.a003707 (2011).

50. Lim, C. & Allada, R. Emerging roles for post-transcriptional regulation in circadian clocks. Nat Neurosci 16, 1544–1550, doi:10.1038/nn.3543 (2013).

51. Cheng, Y., Gvakharia, B. & Hardin, P. E. Two alternatively spliced transcripts from the Drosophila period gene rescue rhythms having different molecular and behavioral characteristics. Mol Cell Biol 18, 6505-6514 (1998).

52. Brunner, M. & Schafmeier, T. Transcriptional and post-transcriptional regulation of the circadian clock of cyanobacteria and Neurospora. Genes Dev 20, 1061–1074, doi:10.1101/gad.1410406 (2006).

53. Colot, H. V., Loros, J. J. & Dunlap, J. C. Temperature-modulated alternative splicing and promoter use in the Circadian clock gene frequency. Mol Biol Cell 16, 5563–5571, doi:10.1091/mbc.E05-08-0756 (2005).

54. Diernfellner, A. C., Schafmeier, T., Merrow, M. W. & Brunner, M. Molecular mechanism of temperature sensing by the circadian clock of Neurospora crassa. Genes Dev 19, 1968–1973, doi:10.1101/gad.345905 (2005).

55. James, A. B. et al. Alternative splicing mediates responses of the Arabidopsis circadian clock to temperature changes. Plant Cell 24, 961–981, doi:10.1105/tpc.111.093948 (2012).

56. Park, M. J., Seo, P. J. & Park, C. M. CCA1 alternative splicing as a way of linking the circadian clock to temperature response in Arabidopsis. Plant Signal Behav 7, 1194–1196, doi:10.4161/psb.21300 (2012).

57. McGlincy, N. J. et al. Regulation of alternative splicing by the circadian clock and food related cues. Genome Biol 13, R54, doi:10.1186/gb-2012-13-6-r54 (2012).

58. Yeoh, L. M., Lee, V. V., McFadden, G. I. & Ralph, S. A. Alternative Splicing in Apicomplexan Parasites. MBio 10, doi:10.1128/mBio.02866-18 (2019).

59. Moller-Levet, C. S. et al. Effects of insufficient sleep on circadian rhythmicity and expression amplitude of the human blood transcriptome. Proc Natl Acad Sci U S A 110, E1132–1141, doi:10.1073/pnas.1217154110 (2013).

60. Etchegaray, J. P., Lee, C., Wade, P. A. & Reppert, S. M. Rhythmic histone acetylation underlies transcription in the mammalian circadian clock. Nature 421, 177–182, doi:10.1038/nature01314 (2003).

61. Jouffe, C. et al. The circadian clock coordinates ribosome biogenesis. PLoS Biol 11, e1001455, doi:10.1371/journal.pbio.1001455 (2013).

62. Koike, N. et al. Transcriptional architecture and chromatin landscape of the core circadian clock in mammals. Science 338, 349–354, doi:10.1126/science.1226339 (2012).

63. Falanga, P. B., D’Imperio Lima, M. R., Coutinho, A. & Pereira da Silva, L. Isotypic pattern of the polyclonal B cell response during primary infection by Plasmodium chabaudi and in immune-protected mice. Eur J Immunol 17, 599–603, doi:10.1002/eji.1830170504 (1987).

64. Ramaprasad, A., Subudhi, A. K., Culleton, R. & Pain, A. A fast and cost-effective microsampling protocol incorporating reduced animal usage for time-series transcriptomics in rodent malaria parasites. Malar J 18, 26, doi:10.1186/s12936-019-2659-4 (2019).

65. Mutungi, J. K., Yahata, K., Sakaguchi, M. & Kaneko, O. Isolation of invasive Plasmodium yoelii merozoites with a long half-life to evaluate invasion dynamics and potential invasion inhibitors. Mol Biochem Parasitol 204, 26–33, doi:10.1016/j.molbiopara.2015.12.003 (2015).

66. Janse, C. J., Ramesar, J. & Waters, A. P. High-efficiency transfection and drug selection of genetically transformed blood stages of the rodent malaria parasite Plasmodium berghei. Nat Protoc 1, 346–356, doi:10.1038/nprot.2006.53 (2006).

67. Oster, H., Damerow, S., Hut, R. A. & Eichele, G. Transcriptional profiling in the adrenal gland reveals circadian regulation of hormone biosynthesis genes and nucleosome assembly genes. J Biol Rhythms 21, 350–361, doi:10.1177/0748730406293053 (2006).

68. Bolger, A. M., Lohse, M. & Usadel, B. Trimmomatic: a flexible trimmer for Illumina sequence data. Bioinformatics 30, 2114–2120, doi:10.1093/bioinformatics/btu170 (2014).

69. Kim, D. et al. TopHat2: accurate alignment of transcriptomes in the presence of insertions, deletions and gene fusions. Genome Biol 14, R36, doi:10.1186/gb-2013-14-4-r36 (2013).

70. Anders, S., Pyl, P. T. & Huber, W. HTSeq--a Python framework to work with high-throughput sequencing data. Bioinformatics 31, 166–169, doi:10.1093/bioinformatics/btu638 (2015).

71. McCarthy, D. J., Chen, Y. & Smyth, G. K. Differential expression analysis of multifactor RNA-Seq experiments with respect to biological variation. Nucleic Acids Res 40, 4288–4297, doi:10.1093/nar/gks042 (2012).

72. Ritchie, M. E. et al. limma powers differential expression analyses for RNA-sequencing and microarray studies. Nucleic Acids Res 43, e47, doi:10.1093/nar/gkv007 (2015).

73. Love, M. I., Huber, W. & Anders, S. Moderated estimation of fold change and dispersion for RNA-seq data with DESeq2. Genome Biol 15, 550, doi:10.1186/s13059-014-0550-8 (2014).

74. Hughes, M. E., Hogenesch, J. B. & Kornacker, K. JTK_CYCLE: an efficient nonparametric algorithm for detecting rhythmic components in genome-scale data sets. J Biol Rhythms 25, 372–380, doi:10.1177/0748730410379711 (2010).

75. Yang, R. & Su, Z. Analyzing circadian expression data by harmonic regression based on autoregressive spectral estimation. Bioinformatics 26, i168–174, doi:10.1093/bioinformatics/btq189 (2010).

76. Wu, G., Anafi, R. C., Hughes, M. E., Kornacker, K. & Hogenesch, J. B. MetaCycle: an integrated R package to evaluate periodicity in large scale data. Bioinformatics 32, 3351–3353, doi:10.1093/bioinformatics/btw405 (2016).

77. Falcon, S. & Gentleman, R. Using GOstats to test gene lists for GO term association. Bioinformatics 23, 257–258, doi:10.1093/bioinformatics/btl567 (2007).

78. Anders, S., Reyes, A. & Huber, W. Detecting differential usage of exons from RNA-seq data. Genome Res 22, 2008–2017, doi:10.1101/gr.133744.111 (2012).

79. Yeoh, L. M. et al. A serine-arginine-rich (SR) splicing factor modulates alternative splicing of over a thousand genes in Toxoplasma gondii. Nucleic Acids Res 43, 4661–4675, doi:10.1093/nar/gkv311 (2015).

