## Supplemental Filles for "Disruption of the coordination between host circadian rhythms and malaria parasite development alters the duration of the intraerythrocytic cycle"

**This document includes:**

Supplementary Fig. 1 to 4
Supplementary Table 1

**Supplementary Information**

Supplementary Data 1. Genes with circadian expression detected in host circadian rhythm matched and mismatched *P. chabaudi* parasites.

Supplementary Data 2. Enriched gene ontology terms associated with host-cues responsive circadian genes.

Supplementary Data 3. Genes with circadian expression detected in *P. falciparum* free running condition.

Supplementary Data 4. Genes with circadian expression detected in *P. chabaudi chabaudi* AS wild-type and *sr10*KO parasites.

Supplementary Data 5. Enriched gene ontology terms associated with SR10 linked circadian genes.

Supplementary Data 6. Differentially regulated genes identified in *sr10*KO parasites compared to wild parasites in 4 matching time points.

Supplementary Data 7. Genes for which differential alternative splicing events detected in two time points of *sr10*KO parasites compared to wild parasites.

Supplementary Data 8. Sequence of primers used in this study.

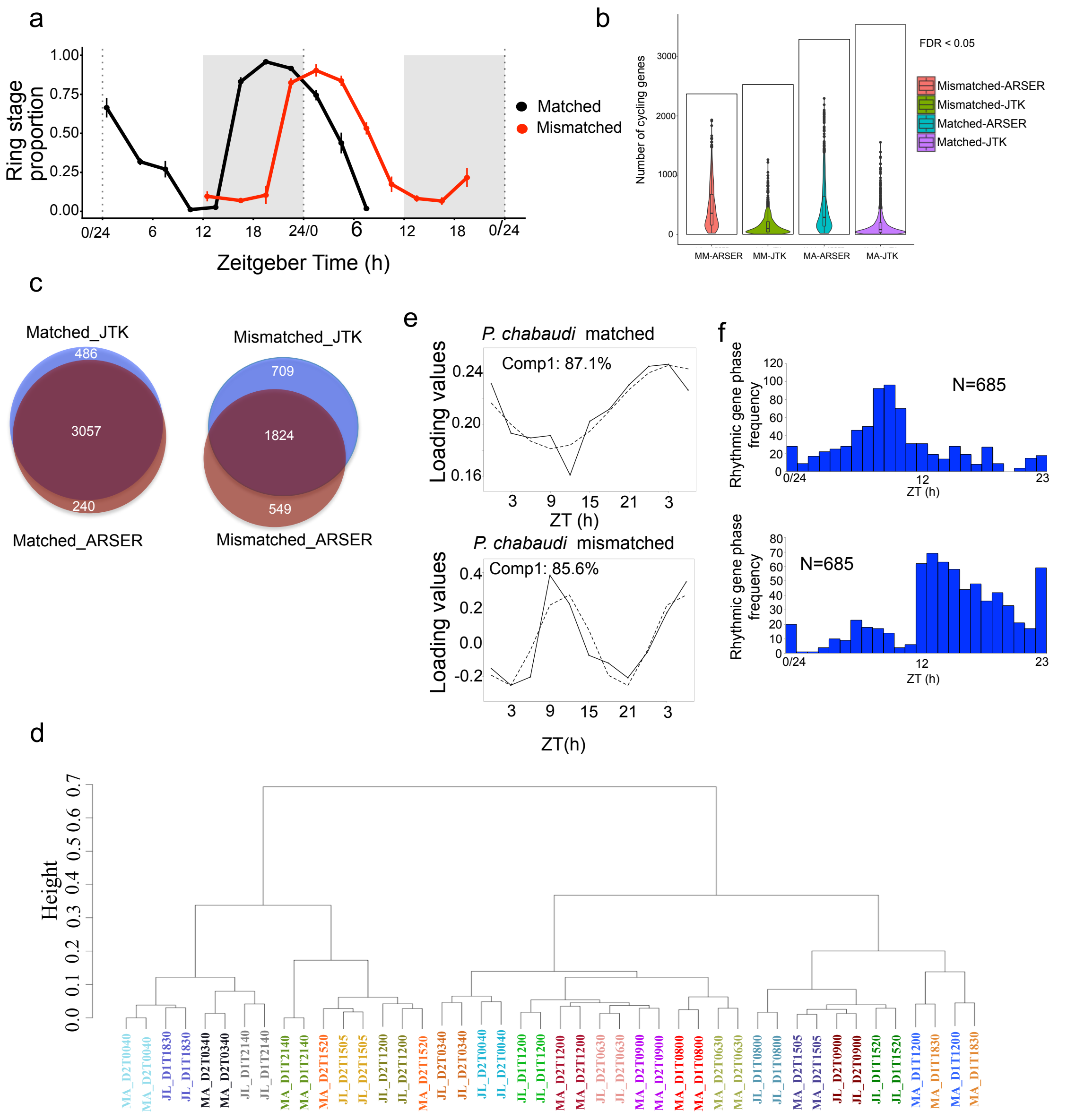


**Supplementary Fig. 1 | Identification of host-cues-responsive circadian transcripts**

a) Ring stage proportion of matched (black line) and mismatched (red line) parasites across 11 time points (Mean ± SEM, N = 4 per time point).

b) Number of circadian genes identified by two algorithms employed in the correct order of sampling time points (black bars). Violin plot represents the distribution of circadian genes identified by permuting the order of sampling time points 1000 times. Box plot inside the violin plots represents the median and quartiles.

c) Venn diagram of the number of circadian transcripts identified by two algorithms (see methods) in matched and mismatched parasites ($p < 0.05$).

d) Clustered dendrogram of 11 time points with two biological replicates per time points using hierarchical clustering algorithm. Analysis was performed on normalized count data Abbreviations: MA, matched; JL, mismatched

e) Principal component analysis (PCA) from matched and mismatched parasites representing cyclic component in solid line and dashed line of its best-fitted cosine curve, which is the first component of the PCA.

f) Rhythmic genes in matched and mismatched parasite that had delayed phase of ~6 h in matched compared to mismatched parasites

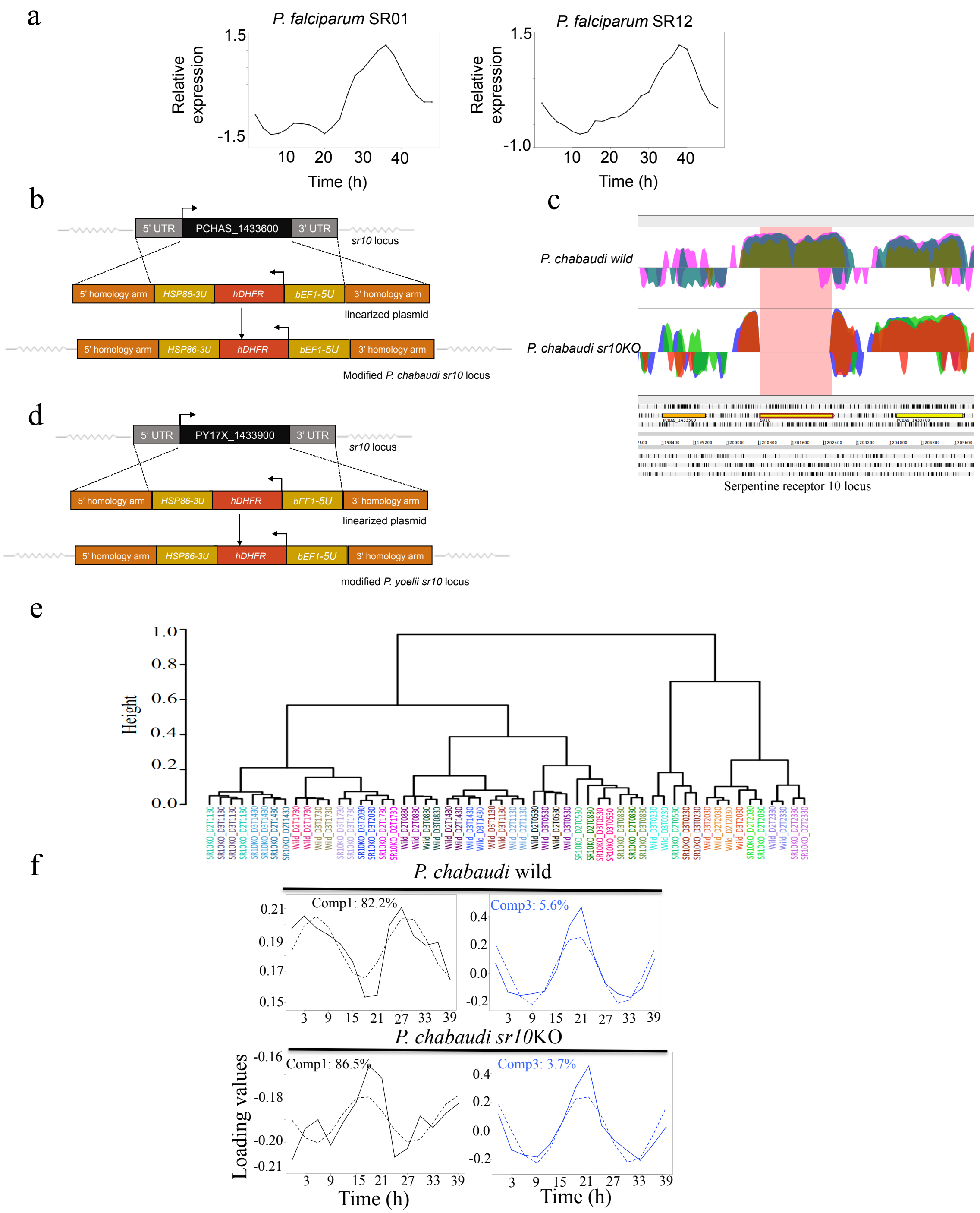


**Supplementary Fig. 2 | Confirmation of *sr10* knockout using RNAseq data.**

a) Expression profiles of *P. falciparum* SR01 and SR12 over 48 h intra-erythrocytic developmental cycle. Microarray data utilized for generating the expression profile was obtained from Bozdech et al., (2003)[1].

b) Schematic of *sr10* knockout strategy in *P. chabaudi*. Linearized plasmid contains a human dihydrofolate reductase-thymidylate synthase (hDHFR-TS) under the control of *P. berghei* EF1 alpha promoter and *P. falciparum* HSP86 3' UTR terminator which is flanked by 1000 bp long regions homologous to 5'UTR and 3'UTR of PCHAS_1433600. Upon integration, the complete PCHAS_1433600 coding region is replaced with the hDHFR drug resistance cassette.

c) Confirmation of *sr10* knockout in *P.chabaudi* using RNAseq. Artemis view of *sr10* locus in *P. chabaudi* wild and *P. chabaudi sr10*KO parasites. *sr10* locus is highlighted. As seen clearly, reads from wild parasites mapped perfectly in the *sr10* region where as no reads from *sr10*KO parasites mapped to the same genomic region. Sequencing reads from 4 randomly chosen time-points from each parasite strains are shown.

d) Schematic of *sr10* knockout strategy in *P. yoelii*. Linearized plasmid contains a human dihydrofolate reductase-thymidylate synthase (hDHFR-TS) under the control of *P. berghei* EF1 alpha promoter and *P. falciparum* HSP86 3' UTR terminator which is flanked by 1000 bp long regions homologous to 5'UTR and 3'UTR of PY17X_1433900. Upon integration, the complete PY17X_1433900 coding region is replaced with the hDHFR drug resistance cassette. Integration and expression (or lack thereof) were confirmed by RNA sequencing.

e) Clustered dendrogram of 14 time points with two biological replicates per time points using hierarchical clustering algorithm. Analysis was performed on normalized count data. The agglomeration algorithm used was ward.D2.

f) Principal component analysis of wild and *sr10*KO parasites representing cyclic in solid line and best fitted cosine curve in dashed line which are first and third components of the PCA.

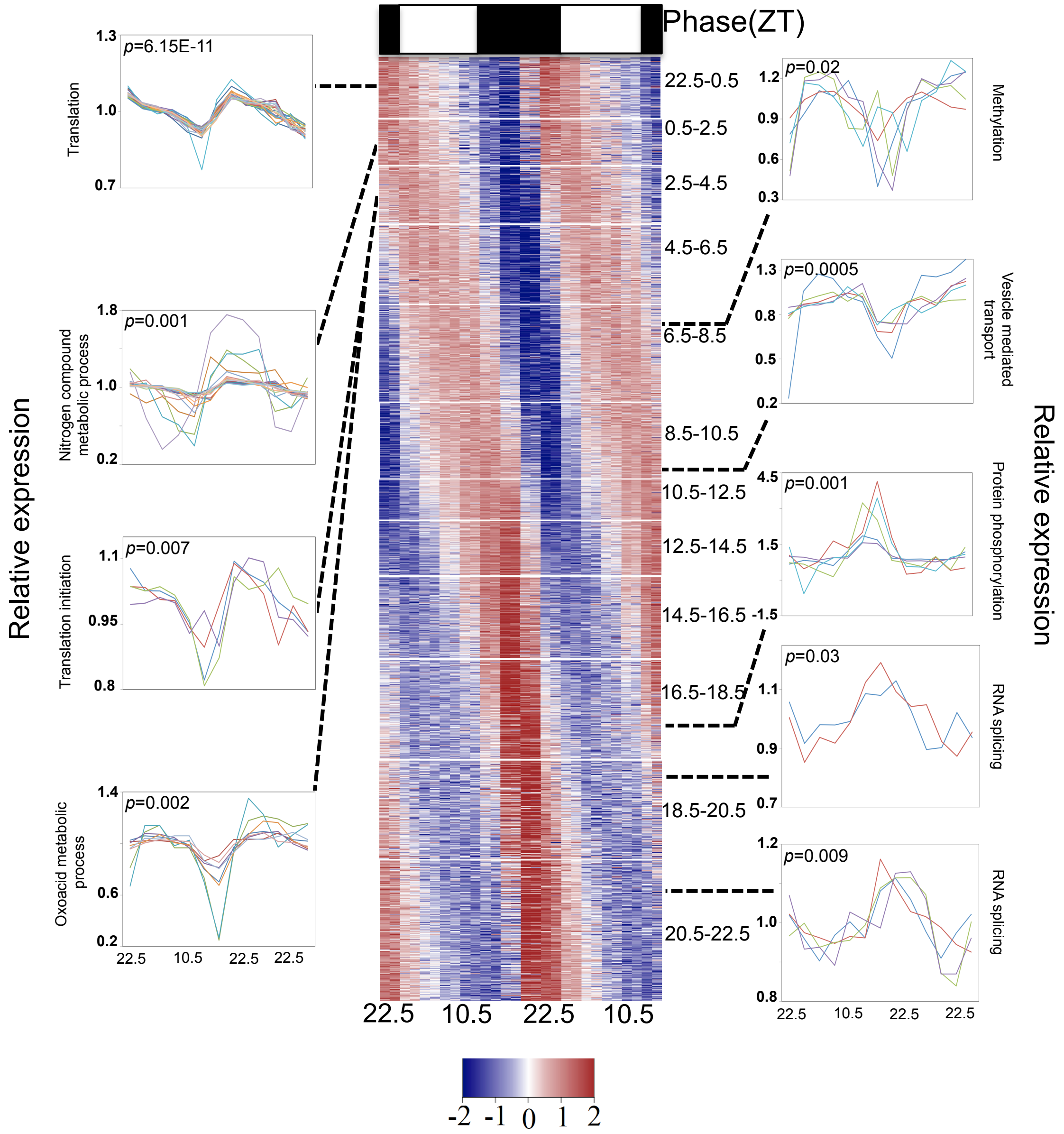


**Supplementary Fig. 3 | Disruption of *sr10* affected circadian genes associated with multiple metabolic processes in *P. chabaudi chabaudi* AS parasites.**

Time series gene expression view of circadian genes that lost rhythmicity in mismatched parasites. The heat map has been segregated into 12 parts with each part representing 2 h phase clusters. Genes were sorted based of phase of expression. Line plots along the sides of the heat map represent expression profiles of individual genes from the significantly enriched gene ontology terms ($p < 0.05$, hypergeometric test) in few selected phase clusters. Each plot has information about the false discovery rate corrected $p$ value of representing gene ontology term. Y axis represents relative expression of genes in each time points which was determined by expression count of each genes normalized by its mean derived from 14 time points.

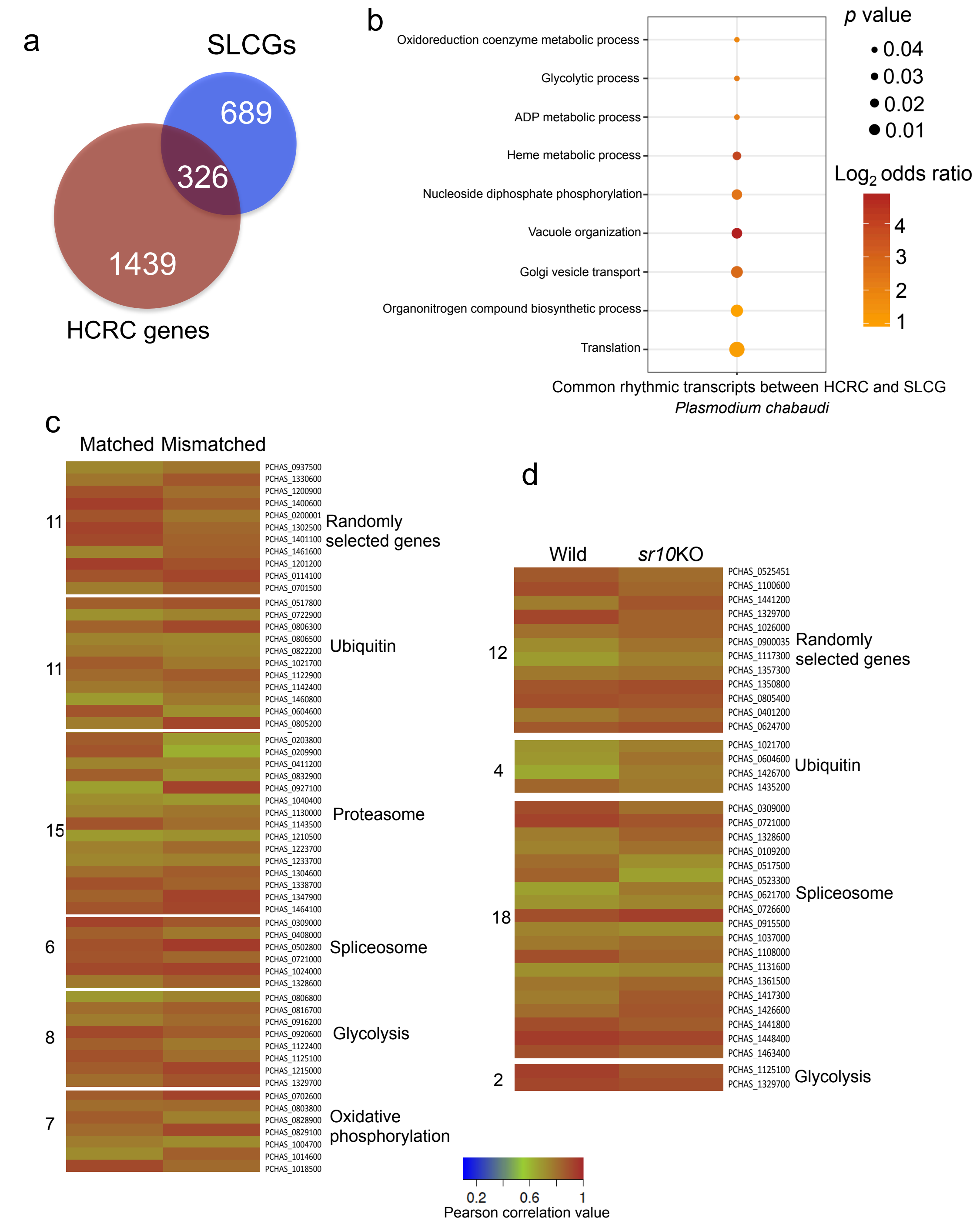


**Supplementary Fig. 4 | Comparison of circadian genes that lost rhythm in mismatched parasites and *sr10*KO parasites.**

a) Venn diagram comparing host responsive circadian genes (HCRC genes) and *sr10*KO linked circadian genes (SLCGs). Circadian genes in matched parasites that lost circadian rhythms in host-rhythm mismatched parasites (HCRC genes) were compared with circadian genes that lost rhythm in *sr10*KO parasites compared to wild-type parasites (SLCGs).

b) Gene ontology enrichment analysis of genes common between host cues responsive circadian genes and SR10 linked circadian genes. Manually curated enriched gene ontology terms ($p$ <0.05, hypergeometric test) have been represented.

c) Heat map representing correlation values between high-throughput qPCR (HT qPCR) and RNASeq data for matched and mismatched parasites. A total of 58 genes were validated.

d) Heat map representing correlation values between HT qPCR and RNASeq data for *P. chabaudi* wild and *P. chabaudi sr10*KO parasites. A total of 36 genes were validated.
RNASeq expression data from 8 time points were compared with HT qPCR expression data from 8 time points to obtain the Pearson correlation values. For RNASeq, normalized count values were used, whereas for qPCR, transcript values (Ct) of genes were normalized to non-cycling transcripts of U5 small nuclear ribonucleoprotein component, putative (PCHAS_1202900). Two technical replicates and two biological replicates were used per time points for HT qPCR experiment. Two biological replicates were used per time points for RNASeq experiment.

Supplementary Table 1: Rhythmicity characteristics of wild type and *sr10*KO parasites.

| Parasite Strain | Parasite Stage | n | Amplitude (mean ± se) | Period (mean ±se) | Phase (mean ±sd) | Mean CoG (mean ±sd) |
|---|---|---|---|---|---|---|
| Pch SR10KO A | early_trophs | 4 | 0.79 ± 0.02 | 22.35 ± 0.43 | 3.19 ± 0.34 | 0.71 ± 0.07 |
| Pch SR10KO B | early_trophs | 4 | 0.93 ± 0.03 | 22.45 ± 0.24 | 1.63 ± 0.16 | 23.45 ± 0.03 |
| Pch WT | early_trophs | 4 | 0.94 ± 0.03 | 25.15 ± 0.37 | 0.16 ± 0.25 | 2.04 ± 0.04 |
| Pyo SR10KO | early_trophs | 4 | 0.30 ± 0.02 | 24.45 ± 0.13 | 20.81 ± 0.11 | 22.37 ± 0.12 |
| Pyo WT | early_trophs | 4 | 0.31 ± 0.01 | 28.03 ± 0.68 | 15.69 ± 0.33 | 20.81 ± 0.25 |


1 Bozdech, Z. *et al.* The transcriptome of the intraerythrocytic developmental cycle of Plasmodium falciparum. *PLoS Biol* **1**, E5, doi:10.1371/journal.pbio.0000005 (2003).
